# The transcription factor ZmbZIP75 promotes both grain filling and kernel dehydration in maize

**DOI:** 10.1101/2024.09.11.612493

**Authors:** Tiandan Long, Yayun Wang, Jin Yang, Zhou Liu, Changqing Mao, Yufeng Hu, Junjie Zhang, Hanmei Liu, Yinghong Liu, Xiujun Fan, Lei Gao, Huanhuan Huang, Ying Xie, Daqiu Zhao, Yubi Huang, Yangping Li

## Abstract

Selecting both high-yield and low-kernel-moisture varieties is essential for modern maize production, but relevant breeding efforts are hindered by a lack of valuable regulatory genes. Here, we demonstrate that the transcription factor (TF) basic leucine zipper 75 (ZmbZIP75) promotes grain yield and reduces kernel moisture in maize. Knockout of ZmbZIP75 results in defective grain filling and kernel dehydration, whereas ZmbZIP75 overexpression confers increased grain yield per plant and decreased kernel moisture without altering plant architecture. Mechanistically, during the grain filling stage, ZmbZIP75 is transcriptionally induced by maternal-derived basal abscisic acid (ABA) and directly activates multiple core starch synthesis-related genes and key TFs, thereby promoting grain filling and final yield. In the late stage of kernel development, high concentrations of zygotic ABA enhance ZmbZIP75 phosphorylation through SnRK2.10. The phosphorylated ZmbZIP75 subsequently transactivates and interacts with TF VP1 to synergistically promote kernel dehydration. This study thus highlights the potential of ZmbZIP75 for engineering both high-yield and low-kernel-moisture varieties to meet the demands of high-efficient maize production.

**IN A NUTSHELL:** *Background:* High grain yield in maize is generally associated with elevated kernel moisture at harvest, which is the main limiting factor for modern maize production. Therefore, it is of importance to select varieties with both high-yield and low-kernel-moisture. These traits are largely controlled by grain filling and kernel dehydration, two tightly connected processes during maize kernel development. Abscisic acid (ABA) is well-documented for its vital role in grain filling and dehydration maturation. However, the molecular mechanisms by which ABA coordinates these two processes remain unclear.

*Question:* What are the regulatory factors involved, and how do they mediate ABA signaling to coordinate grain filling and dehydration maturation in maize?

*Findings:* ZmbZIP75 is transcriptionally induced by ABA and directly activates multiple core starch synthesis-related genes and key TFs in developing maize endosperm, thereby promoting grain filling. In developing embryo, ABA enhances ZmbZIP75 phosphorylation via SnRK2.10. The phosphorylated ZmbZIP75 then transactivates and interacts with VP1 to synergistically promote kernel dehydration. Moreover, *ZmbZIP75* overexpression confers increased grain yield and reduced kernel moisture in maize.

*Next steps:* While ZmbZIP75 is directly phosphorylated to mediate ABA signaling during dehydration maturation, *ZmbZIP75* is transcriptionally induced by ABA during grain filling. We plan to identify upstream factors that mediate ABA signaling to regulate *ZmbZIP75* expression, thereby enhancing our understanding of ABA-promoted grain filling in maize.

## Introduction

Global maize production needs to be constantly increased with high efficiency to meet the rising demand for human food and livestock products, owing to the growing population and rapid economic development (Hickey et al. 2019; Erenstein et al. 2022). Over the past decades, maize breeders have tended to select late-maturing varieties to pursue higher grain yield through maximizing regional thermal resources, especially in developing countries such as China (Tao et al. 2014; Massigoge et al. 2023). However, maize high grain yield is accompanied by high kernel moisture at harvest, which is the main limiting factor for mechanized grain harvesting (Chai et al. 2017). A high kernel moisture at harvest also results in high drying cost and grain mildew rate that directly impairs maize quality (Xie et al. 2022). Therefore, it is essential to ameliorate current grain yield-moisture paradigm by breeding new varieties with both high grain yield and low kernel moisture for further maize production.

Maize kernel development is generally divided into three partially overlapping phases (Zhang and Kaeppler 2017b). The initial lag phase is characterized by active cell proliferation and differentiation, exhibiting a dramatic increase in moisture content with minimal dry matter accumulation. The subsequent grain filling phase involves the rapid accumulation of storage materials, while kernel moisture content gradually declines. In the last post-maturity dry-down phase, kernels reach maximum weight, coupled with a steeper decline in moisture content and subsequently enter a quiescent state (Maiorano et al. 2014). Two grain filling-related traits, grain filling rate and grain filling duration, determine the final grain yield (Johnson and Tanner 1972; Gasura et al. 2013; Xing et al. 2023). On the other hand, the kernel dehydration rate largely determines the kernel moisture content at harvest (Liu et al. 2020). Thus, in order to achieve high-yield and low-kernel-moisture varieties, it is necessary to accelerate the grain filling rate within a sufficient grain filling duration while maximizing the kernel dehydration rate (Xu et al. 2022).

Maize grain filling involves the transport of assimilates to grains via the basal endosperm transfer layer (BETL) and the accumulation of reserve materials such as starch and storage protein in the endosperm, as well as lipids in the embryo (Ma et al. 2023). Starch is the major determinant for maize yield because it accounts for approximately 70% of grain dry weight (Hannah and James 2008). Starch biosynthesis requires the synergistic activity of multiple key enzymes, including ADP-glucose pyrophosphorylase (AGPase), starch synthase (SS), starch branching enzyme (SBE), and starch debranching enzyme (DBE) (James et al. 2003; Huang et al. 2021). Each class of these enzymes is encoded by multiple genes in various maize tissues (Yan et al. 2009). Previous transcriptome analyses have demonstrated that core starch synthesis-related genes (SSRGs) in maize endosperm share a similar expression pattern during grain filling, indicating that these genes may be tightly co-regulated by specific transcription factors (TFs) (Prioul et al. 2008; Chen et al. 2014; Hu et al. 2021). In recent years, several TFs involved in starch synthesis have been characterized in maize endosperm, including NKD1/2 (Gontarek et al. 2016), Opaque 2 (Zhang et al. 2016; Deng et al. 2020), and ZmNAC128/130 (Zhang et al. 2019b; Chen et al. 2023), these TFs directly regulate the expression of several SSRGs, and mutations in these TF genes impair endosperm starch accumulation and grain filling.

Upon the termination of grain filling, maize kernels initiate the process of dehydration and dormancy. Numerous proteins associated with seed maturation and desiccation tolerance, such as LEA (late embryogenesis abundant) proteins and HSPs (heat shock proteins), are synthesized at a high rate during this stage (Amara et al. 2011; Zhang et al. 2024). The expression of these genes is predominantly controlled by the phytohormone abscisic acid (ABA) (Wu et al. 2015; Niu et al. 2022). ABA plays a central role in the regulation of kernel dehydration process. White maize typically has higher kernel moisture at harvest compared to yellow maize (Kang and Zuber 1989), due to the absence of phytoene synthase, an enzyme essential for carotenoid and ABA synthesis within the kernels (Buckner et al. 1996). Likewise, maize mutants with disruptions in *Vp2*, *Vp5*, and *Vp7*, which block ABA synthesis in the kernel, display significantly higher kernel moisture content than the corresponding wild-types (Wilson et al. 1973; Neill et al. 1986). Additionally, ABA-mediated kernel dehydration and dormancy depends on the presence of a classic B3 domain-containing TF, VIVIPAROUS1 (VP1). *vp1* mutants exhibit insensitivity to ABA, leading to unsuccessful kernel dehydration and dormancy (Neill et al. 1986; McCarty et al. 1989).

In addition to its central function in maturation drying process, ABA also plays a vital role in the regulation of crop grain filling. Large-grain rice variety exhibit higher grain ABA content and faster grain-filling rate compared to the small-grain variety (Kato et al. 1993). Under moderate soil drought conditions, the increased ABA content in developing grains enhances the activities of starch synthetic enzymes, thereby promoting the grain-filling rate and grain weight (Yang et al. 2004; Zhang et al. 2012; Yuan et al. 2023). Exogenous ABA application also significantly accelerates grain filling through enhancing endosperm starch synthesis in rice and maize kernel (Yang et al. 2001; Yu et al. 2024). Rice DG1 transporter and its maize homolog mediate the long-distance transport of ABA from maternal tissues to grains, and mutation of this gene significantly reduces grain ABA content, hampering starch synthesis and grain filling (Qin et al. 2021). Surprisingly, maize *vp2*, *vp5*, and *vp7* mutants with defective kernel ABA synthesis exhibit almost normal starch synthesis and grain filling (Ober and Setter 1992; Wang et al. 2021). These findings indicate that maize grain filling and dehydration are controlled by distinct ABA signaling originating from different tissues. Maternal plant-derived ABA primarily governs grain filling, whereas ABA synthetized *in situ* in the kernel is essential for grain dehydration. Notably, grain filling and kernel dehydration are two closely related stages during maize kernel development. It has been observed that maize kernels with elevated ABA levels exhibit both accelerated grain filling and kernel dehydration (Zhang et al. 2018). However, the mechanism by which ABA coordinates these two processes in maize remain poorly understood.

In this study, we identified a kernel-preferential TF, ZmbZIP75, that plays a dual role in grain filling and kernel dehydration in maize. ZmbZIP75 was induced by ABA and exhibited differential phosphorylation in the endosperm and embryo. Knockout of *ZmbZIP75* impaired grain filling and delayed kernel dehydration, resulting in smaller kernels with higher moisture content at harvest. ZmbZIP75 directly regulates the expression of core SSRGs, late embryo maturation-related genes, and several key TF genes, indicating its central regulatory role in grain filling and kernel dehydration processes. Notably, transgenic lines overexpressing *ZmbZIP75* and the resulting F1 hybrids exhibited increased grain yield and reduced kernel moisture content without affecting normal plant growth. This study thus provides a potential target for breeding high-yield and low-kernel-moisture maize varieties.

## Results

### ABA promotes grain filling and starch synthesis

To identify the regulatory factors responsible for ABA-promoted grain filling, we first investigated the effect of different concentrations of ABA on kernel development in maize. 10 days after pollination (DAP) maize kernels with uniform size were detached and incubated on semisolid half-strength MS medium supplemented with varying ABA concentrations (Supplemental Figure S1). Under 100 μM ABA treatment, maize kernels exhibited accelerated growth and increased grain plumpness and grain weight (Figure 1A and 1B). Iodine staining and starch content analysis revealed that the 100 μM ABA treatment significantly enhanced endosperm starch accumulation (Figure 1C and 1D). In contrast, application of a high ABA concentration (300 μM) did not result in a significant increase in kernel weight or starch content (Figure 1B-1D). These results suggest that a moderate increase in ABA promotes maize grain filling and starch synthesis.

**Figure 1.**
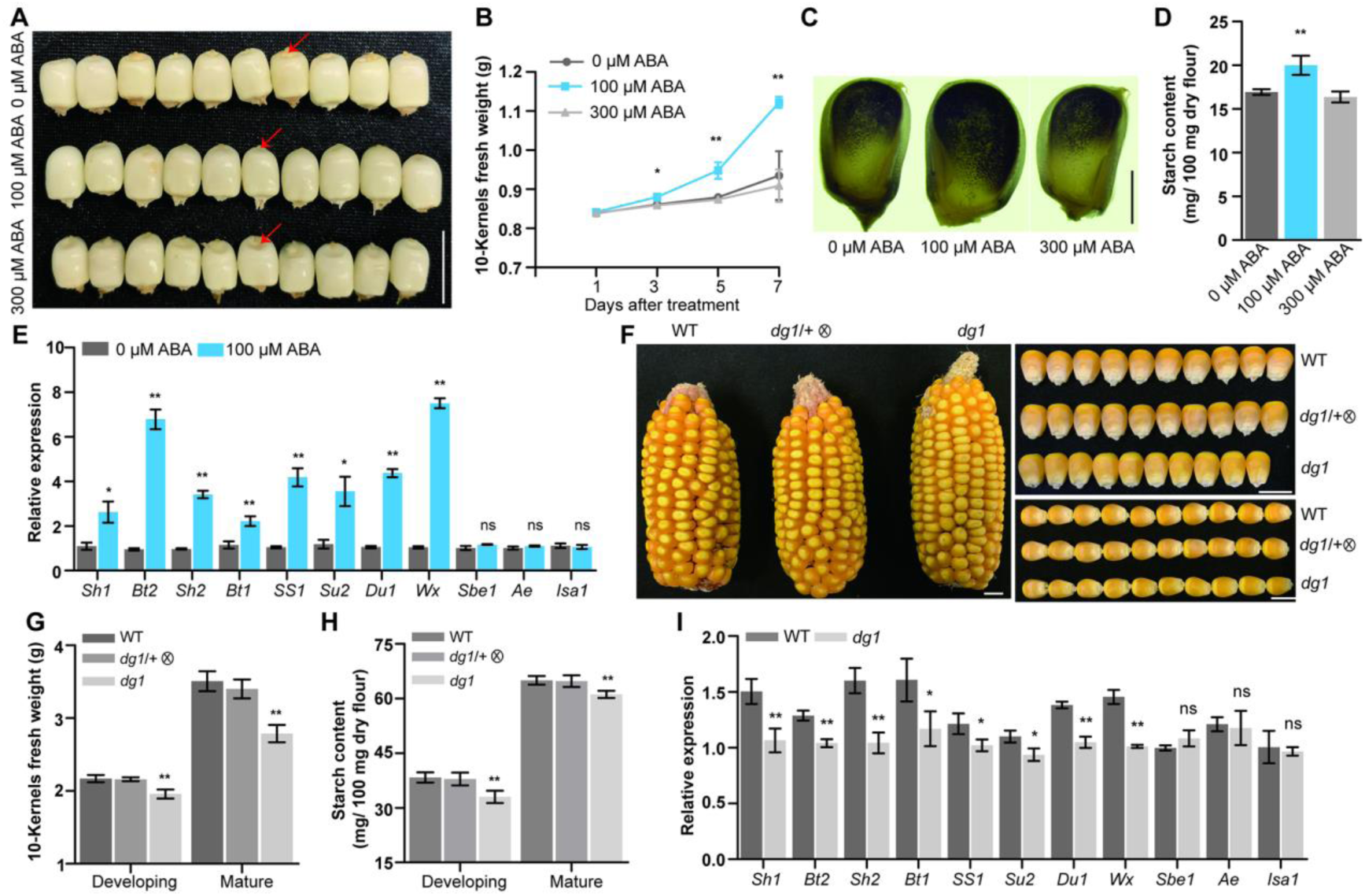
ABA enhances starch synthesis in developing maize kernels. (A) The phenotype of the developing kernels treated with different concentrations of ABA after 5 days. Scale bar, 1 cm. (B) Comparisons of 10-kernels fresh weight of developing kernels treated with 0 μM, 100 μM or 300 μM ABA. Error bars represent SD from three biological replicates. (C) The visualization of the starch in maize kernels using I_2_ /KI staining with 0 μM, 100 μM or 300 μM ABA treatment. Scale bar, 2 mm. (D) Comparisons of the starch content of maize kernels treated with 0 μM, 100 μM or 300 μM ABA. Error bars represent SD from three biological replicates. (E) RT-qPCR analysis of these core starch synthesis-related genes (SSRGs) expression in maize kernels treated with 0 μM or 100 μM ABA treatment. All expression levels were normalized to *ZmActin*. Error bars represent means±SD from three biological replicates. (F) The ear phenotype of self-pollinated WT, *dg1*/+, *dg1* and the observations of kernel size. Scale bar, 1 cm. (G) and (H) Measurement and comparisons of the 10-kernels weight (G) and starch content (H) of self-pollinated WT, *dg1*/+, *dg1* kernels at developing period (18-DAP) and maturation period (36-DAP). (I) RT-qPCR analysis of these core SSRGs expression in WT and *dg1* kernels at 18-DAP. All expression levels were normalized to *ZmActin*. Error bars represent SD from three biological replicates. Statistical significance (*\*P < 0.05*; *\*\*P < 0.01*) was determined by two-tailed Student’s *t*-test.

Starch synthesis in maize kernel is primarily controlled by a set of endosperm-specific core SSRGs (Hu et al. 2021). To determine whether these genes are ABA-inducible, reverse transcription quantitative PCR (RT-qPCR) analysis was performed. The results showed that 8 out of 11 SSRGs were significantly induced by 100 μM ABA, including *Sh1* (sucrose synthase, SUS), *Sh2* and *Bt2* (AGPase large subunit and small subunit, respectively), *Bt1* (ADPG transporter), *SS1* (starch synthase I), *Su2* (starch synthase IIa), *Du1* (starch synthase IIIa), and *Wx* (granule-bound starch synthase I, GBSSI) (Figure 1E).

Maize *DG1* encodes a rice homologous MATE transporter that mediates long-distance ABA transport to developing grains (Qin et al. 2021). To confirm the essential role of ABA in maize grain filling and starch synthesis, an EMS mutant of *DG1* was obtained in the B73 background, which harbors a premature stop codon in the second exon (Lu et al. 2018) (Supplemental Figure S2A). Mutation of *DG1* leads to a significant decrease in ABA content during the rapid grain filling stage (18 DAP) (Supplemental Figure S2B). Consequently, the mutant kernels exhibited a smaller grain size, significantly decreased grain weight, and reduced starch content (Figure 1F-1H). Furthermore, RT-qPCR analysis revealed that the expression of most endosperm-specific core SSRGs was down-regulated in the *dg1* kernels compared to the wild-type (Figure 1I), which is in contrast to the results observed in the kernels treated with 100 μM ABA, where the majority of these SSRGs were significantly induced (Figure 1E). Collectively, these results demonstrate that ABA enhances starch synthesis in maize kernels by activating the expression of core SSRGs, which in turn promotes grain filling.

### Identification of candidate TFs that mediates ABA-inducible grain filling

The coordinated induction of most core SSRGs in maize kernels by ABA (Figure 1E) suggests that common *cis*-elements may exist in their promoters to mediate ABA signaling. To investigate this, we retrieved the 1.5-kb promoter sequences upstream of the start codon (ATG) of eight ABA-inducible SSRGs and conducted motif enrichment analyses using MEME (https://meme-suite.org/meme/tools/meme) (Bailey et al. 2015). Among the top four significantly enriched motifs, Motif 2 was found to be a typical ACGT-containing ABA-responsive element (ABRE) (Supplemental Figure S3A). Further sequence analysis revealed that multiple ABREs are present in the promoters of these eight SSRGs (Supplemental Figure S3B).

The ABRE element has been widely reported to mediate ABA signaling through direct interaction of A-group bZIP TFs (Cutler et al. 2010; Wang et al. 2019a). Using the amino acid sequences of 13 A-group bZIP TFs from Arabidopsis as queries (Dröge-Laser et al. 2018), we performed a BLAST search and identified 16 putative A-group bZIP TFs in the maize reference genome (Zm-B73-REFERENCE-GRAMENE-4.0) (Supplemental Data Set S1). To investigate the role of these TFs in grain filling, we resequenced the genomic fragments (from -1500 bp upstream of the transcription start site (TSS) to the stop code) of these 16 genes from 204 maize inbred lines (Supplemental Data Set S2), and identified a total of 1851 single nucleotide polymorphisms (SNPs) with minor allele frequency (MAF) >5%, missing rate < 5%, and heterozygosity < 20%. Given the tight correlation between grain filling and final grain weight, we conducted association analysis of all these SNPs with 100-grain weight (HGW) using the rMVP package (version 1.0.6) in R (version 4.2.2). Notably, three adjacent SNPs on chromosome 8 were found to be significantly associated with HGW (−log_10_(*P*)>4.0), and the gene underlying this association is *ZmbZIP75* (Zm00001d012296) (Figure 2A). Our previous study demonstrated that *ZmbZIP75* is one of 47 candidate TFs involved in sucrose and ABA-induced expression of SSRGs in maize endosperm (Huang et al. 2016). We further investigated the association of natural variations in these 47 TFs with HGW. A total of 6421 SNPs were identified by resequencing and subjected to association analysis. The results showed that multiple SNPs located in *ZmbZIP75* exhibited stronger associations with HGW compared to those located in the remaining 46 TFs (Figure 2B).

**Figure 2.**
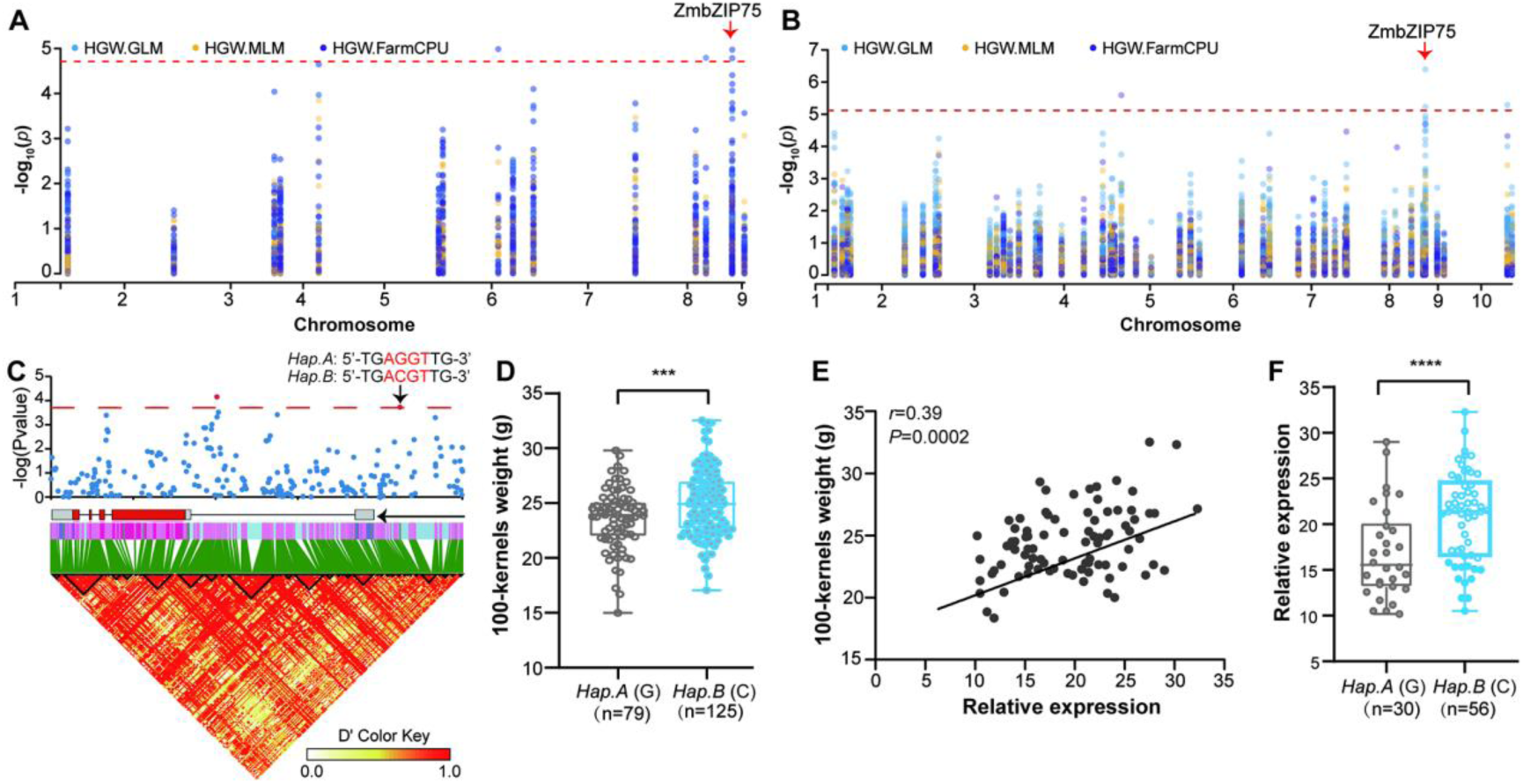
*ZmbZIP75* is a candidate gene for the regulation of ABA-inducible grain filling. (A) Association analysis of group-A bZIP transcription factors that may be involved in ABA regulating kernel filling with HGW-best linear unbiased estimate (HGW-BLUE). (B) Association analysis of 47 candidate transcription factors that may be involved in ABA regulating kernel filling with HGW-BLUE. (C) Association of SNPs within a -1500 bp region of the ZmbZIP75 variations with 100-kernels weight. The significant association loci were highlighted by a red triangle. (D) Boxplot showing the 100-kernel weight of *Hap. A* (G, n=79) and *Hap. B* (C, n=125). The box shows median and interquartile range, and whiskers extend to maximum and minimum values. (E) Pearson correlation between ZmbZIP75 expression and the 100-kernels weight among 86 inbred lines. (F) Boxplot showing the *ZmbZIP75* expression among 86 diverse inbred lines of the *HapA* (G, n=30) and *HapB* (C, n=56). Statistical significance (*\*P < 0.05*; *\*\*P < 0.01*) was determined by two-tailed Student’s *t*-test.

Gene resequencing analysis revealed that 285 SNPs were located in *ZmbZIP75* (Supplemental Data Set S3). We investigated whether these SNPs contribute to the variations in HGW through association analysis. One SNP (SNP250) located in the promoter region was found to be significantly associated with HGW. Based on the variation at SNP250, two haplotypes (Hap.A and Hap.B) were classified among 204 maize inbred lines (Figure 2C), at which Hap.A (SNP250-G) and Hap.B (SNP250-C) were associated with lower and higher HGW, respectively (Figure 2D). Interestingly, SNP250 is located within the putative ABRE motif of *ZmbZIP75* promoter (Figure 2C). We therefore test whether *ZmbZIP75* is ABA-inducible. RT-qPCR showed that *ZmbZIP75* was significantly induced by different concentrations of ABA (50-150 μM) in developing kernels, with maximum induction at 100 μM. While treatment with a high concentration of ABA (300 μM) did not show significant induction effect (Supplemental Figure S4A and 4B). Conversely, *ZmbZIP75* transcription was significantly reduced in the *dg1* kernels compared to the wild-type (Supplemental Figure S4C). This ABA-responsive regulation of *ZmbZIP75* transcription is consistent with the observed effects of ABA on grain filling and starch synthesis (Figure 1).

We next analyzed the association between *ZmbZIP75* transcription and HGW. The expression data of *ZmbZIP75* in 86 maize inbred lines were obtained from PPRD database (https://plantrnadb.com/zmrna/) (Yu et al. 2022) (Supplemental Data Set S4). The result indicated that *ZmbZIP75* transcription was positively correlated with HGW (*r* = 0.39, *P* = 0.0002, Figure 2E), and inbred lines carrying *Hap.B (C)* exhibited higher *ZmbZIP75* expression than those carrying *Hap.A (G)* (with a mutated ABRE motif) (Figure 2F). In addition, preliminary expression analysis indicated that among all 16 A-group bZIP TFs, only *ZmbZIP75* is predominantly expressed in developing maize kernel based on the released transcriptome data (Chen et al. 2014) (Supplemental Figure S5). Taken together, these findings suggest that *ZmbZIP75* may play a regulatory role in mediating ABA-promoted grain filling.

### ZmbZIP75 is kernel-predominantly expressed with differential phosphorylation in endosperm and embryo

Phylogenetic analysis was performed using the full-length amino acid sequences of A-group bZIP TFs from maize, rice and Arabidopsis (Supplemental Figure S6A). ZmbZIP75 was closely related to ABI5 homologs in rice (OsABI5) and Arabidopsis (AtABI5). Sequence alignment analysis revealed that these three bZIP proteins contain four highly conserved domains (M1, M2, M3, and bZIP). The overall amino acid similarity between ZmbZIP75 and OsABI5 was 63.44%, while ZmbZIP75 shared only 37.25% similarity with AtABI5 (Supplemental Figure S6B). We further constructed a phylogenetic tree using ZmbZIP75 and its homologs from other species. The result indicated that ZmbZIP75 is highly conserved in monocots (Supplemental Figure S6C).

We next examined the expression pattern of *ZmbZIP75* in various tissues by RT-qPCR and found that *ZmbZIP75* is predominantly expressed in developing kernels, with expression starting at 6 DAP, gradually increasing throughout kernel development, and peaking at 30 DAP (Figure 3A). Western blot analysis demonstrated that ZmbZIP75 protein accumulation exhibited a similar temporal pattern as the transcript level (Figure 3B). The spatial distribution of *ZmbZIP75* within maize kernel was examined by RNA *in situ* hybridization. The results showed that *ZmbZIP75* is expressed in most compartments of the 15 DAP kernel, with strong signal detected in the embryo (Em) and relatively lower signal in the starchy endosperm (SE). *ZmbZIP75* expression was also observed in the aleurone (AL) and the BETL (Figure 3C). Further immunofluorescence analysis confirmed the spatial distribution of ZmbZIP75 within the 15 DAP kernel at the protein level (Supplemental Figure S7). These results suggest a regulatory role for ZmbZIP75 in both embryo and endosperm development.

**Figure 3.**
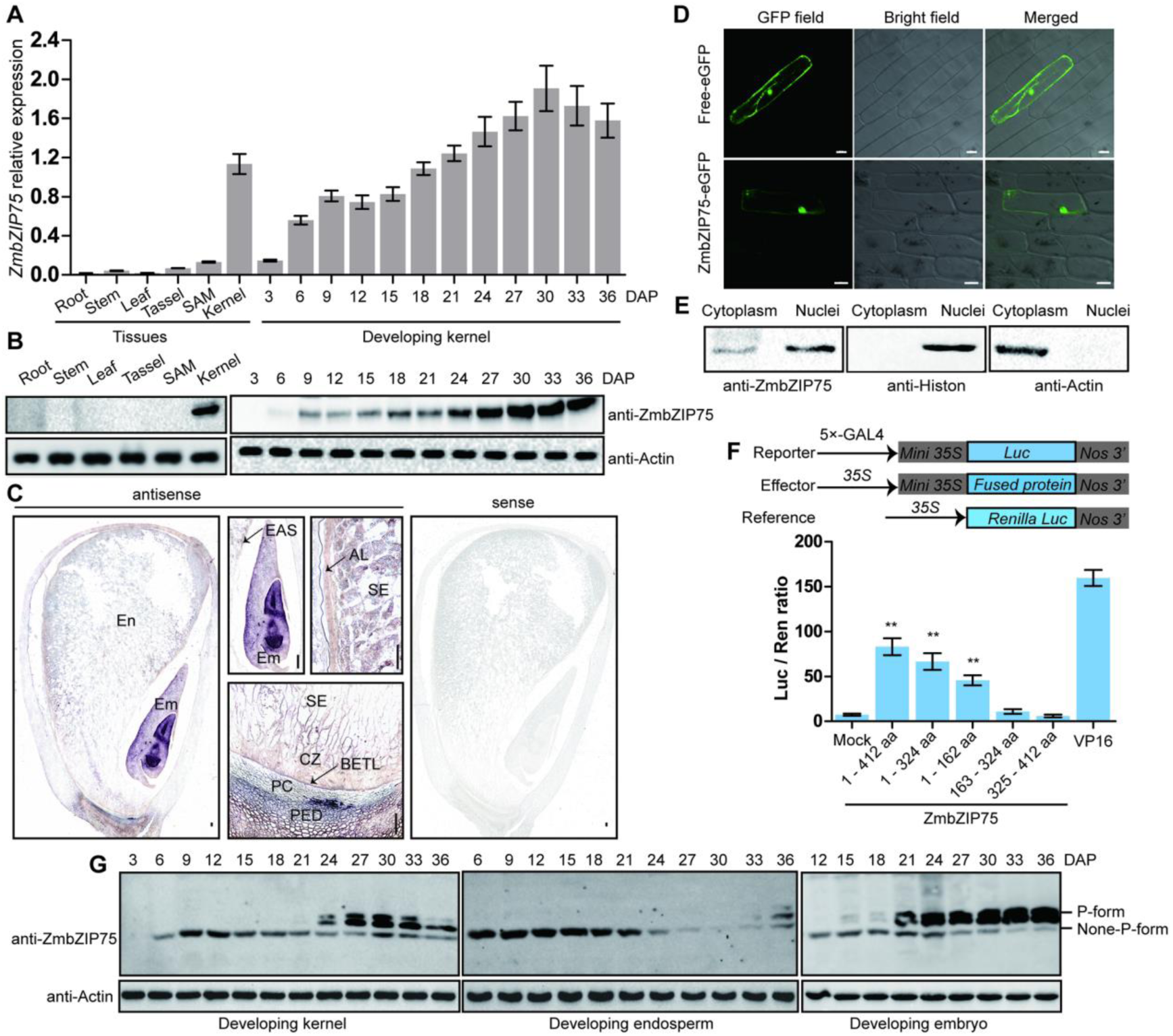
ZmbZIP75 is a kernel-preferential transcription factor. (A) RT-qPCR analysis of *ZmbZIP75* expression in different tissues and developing kernels. All expression levels were normalized to *ZmActin*. Error bars represent means±SD from three biological replicates. SAM, shoot apical meristem. DAP, days after pollination. (B) Immunoblot analysis of ZmbZIP75 in different tissues and developing kernels. Actin was used as a loading control. Scale bar, 50 μm. (C) RNA *in situ* hybridization of ZmbZIP75. Longitudinal sections of 15-DAP kernel were hybridized with an antisense ZmbZIP75 probe. En, endosperm; Em, embryo; EAS, endosperm adjacent to scutellum; AL, aleurone; SE, starchy endosperm; CZ, conducting zone; PC, placento-chalazal region; PED, the vascular region of the pedicel; BETL, the basal endosperm transfer layer. Scale bar, 200 μm. (D) Subcellular localization of ZmbZIP75-eGFP in onion epidermal cells. Free-eGFP was used as a control. Scale bar, 20 μm. (E) Immunoblot analysis of nuclear and cytoplasmic fraction proteins obtained from 9-DAP B73 maize kernels with antibodies against ZmbZIP75, histone H3 (nuclear marker) and Actin (cytoplasmic marker). (F) Transactivation analysis of the full-length ZmbZIP75 and its different fragments in maize leaf protoplasts. Error bars represent SD from three biological replicates. Mock means reporter and reference plasmids only, GALl4-VP16 was used as a positive control. (G) Phosphorylation of ZmbZIP75 in developing kernel, endosperm and embryo, respectively. Total proteins extracted from developing kernels were separated in a Phos-Tag gel, and the phosphorylated (P-form) or nonphosphorylated (None-P-form) ZmbZIP75 proteins were detected with anti-ZmbZIP75 antibody. Actin was used as a loading control. Statistical significance (*\*P < 0.05*; *\*\*P < 0.01*) was determined by two-tailed Student’s *t*-test.

To investigate the subcellular localization of ZmbZIP75, its full-length ORF was fused to enhanced GFP (eGFP) and driven by constitutive *35S* promoter. The fusion construct was transformed into onion epidermal cells by particle bombardment. Imaging of the bombarded epidermal cells revealed that the ZmbZIP75-eGFP fusion protein was localized primarily in the nucleus, with some additional localization observed in the cytoplasm (Figure 3D). To further confirm the subcellular distribution of ZmbZIP75, nuclear and cytoplasmic protein fractions were isolated from 9 DAP maize kernels and subjected to immunoblot analysis. The results also showed that ZmbZIP75 exhibited both nuclear and cytosolic localization (Figure 3E).

The transcriptional activity of ZmbZIP75 was assessed using a dual-luciferase reporter system. The full-length *ZmbZIP75*, as well as a series of truncated fragments, were fused to the yeast GAL4 DNA-binding domain (GAL4-BD) as effectors. These effector constructs were then co-transformed into maize protoplasts, along with a reporter construct containing a luciferase (LUC) gene driven by a promoter with five copies of the GAL4 binding site. The results revealed that the full-length ZmbZIP75 possesses strong transcriptional activation activity. However, deletion of the N-terminal 1-162 amino acids abolished this transactivation function (Figure 3F). Consistent results were also observed in yeast transactivation assays (Supplemental Figure S8), suggesting that this N-terminal region is essential for the transactivation activity of ZmbZIP75.

The N-terminal region (1-162 amino acids) of ZmbZIP75 contains two conserved domains, M1 and M2, each harboring a putative phosphorylation recognition motif (R-X-X-S) (Supplemental Figure S9A). To examine whether ZmbZIP75 is phosphorylated *in vivo*, we immunoprecipitated ZmbZIP75 from *ZmbZIP75-Flag* overexpressing kernels at 18 DAP with anti-Flag antibody. Immunoblotting with an anti-phosphoserine antibody detected a phosphorylation signal, which was eliminated upon treatment with λPPase (Supplemental Figure S9B). Subsequently, we detected the phosphorylation dynamics of ZmbZIP75 during kernel development using Phos-tag SDS-PAGE (Figure 3G). Phosphorylated ZmbZIP75 in developing kernels accumulate by 24 DAP and peaks by 30 DAP. When the endosperm and embryo were separated after 12 DAP for Phos-Tag analysis, non-phosphorylated ZmbZIP75 was detected in the endosperm throughout the grain filling stage. In contrast, ZmbZIP75 was predominantly found in a phosphorylated state in the embryo, especially during the late stages of grain maturation. These findings suggest that the distinct phosphorylation states of ZmbZIP75 in the endosperm and embryo may underlie its divergent regulatory functions during maize grain filling and late grain maturation.

### Knockout of *ZmbZIP75* causes defective grain filling and delayed grain dehydration

To investigate the physiological roles of *ZmbZIP75*, knockout mutants were generated in the KN5585 inbred line using CRISPR-Cas9 system. Two independent knockout lines, *zmbzip75-15* and *zmbzip75-18*, were identified with a 1-bp deletion at positions +249 and +245 relative to the ATG, respectively, leading to frameshifts and premature stop codons (Supplemental Figure S10A). Immunoblot analysis confirmed the complete absence of the ZmbZIP75 protein in both knockout lines (Supplemental Figure S10B). These two mutants were crossed with the wild-type, and then the Cas9-free plants were self-pollinated. Due to the poor germination of the mutant kernels, the heterozygous F2 ears were selected for further analysis (Supplemental Figure S10C). The representative F2 ears (*zmbzip75-15/+* and *zmbzip75-18/+*) exhibited a 3:1 segregation ratio of wild-type and mutant kernels (Figure 4A). Compared to the wild-type, the *zmbzip75* kernels were significantly smaller, with reduced kernel width and thickness (Figure 4B; Supplemental Figure S10D-10F). The kernel weight of the mutants in the *zmbzip75-15/+* and *zmbzip75-18/+* ears was only 45% and 34% of the corresponding wild-type, respectively, indicating a defect in grain filling (Figure 4C). Further analysis of the dynamic dry matter accumulation revealed that the *zmbzip75* kernels had a significantly lower grain dry weight and grain filling rate than the wild-type throughout kernel development (Figure 4D and 4E).

**Figure 4.**
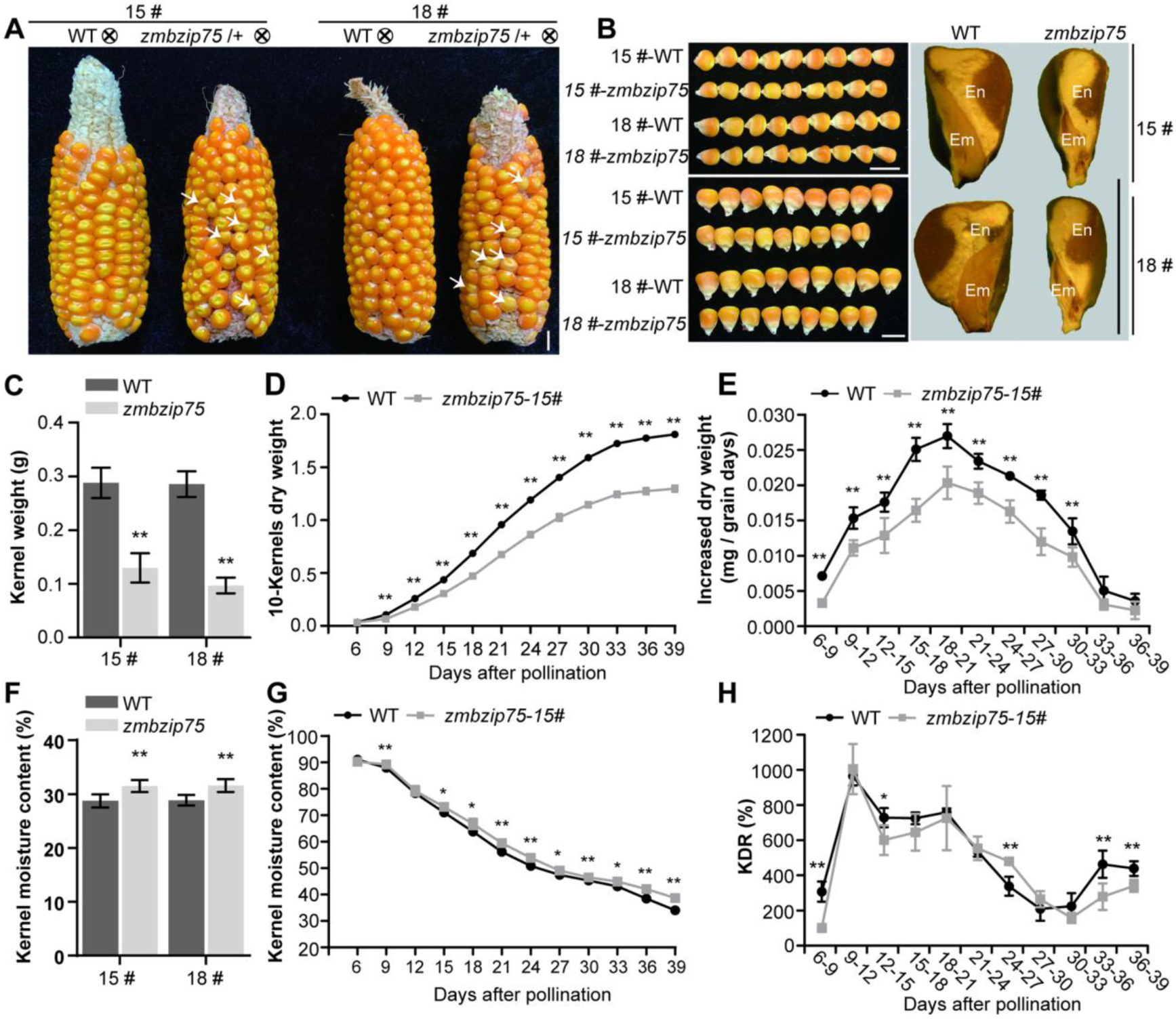
Phenotypic analysis of *zmbzip75* kernels. (A) The ear phenotype of self-pollinated *zmbzip75*-15-/+, *zmbzip75*-18-/+ and WT of two mutant lines. White arrow indicated *zmbzip75* mutant kernels. Scale bar, 1 cm. (B) The observations of kernel size of WT and *zmbzip75* mutants. Left panel, comparisons of the kernel length and width of WT and *zmbzip75*. Right panel, comparisons of the kernel thickness between WT and *zmbzip75* through a longitudinal section. Scale bar, 1 cm. (C) Measurement and comparisons of kernel weight of WT and *zmbzip75*. Data were means± SD (*n* = 20 biologically independent samples). (D) and (E) Measurement and comparisons of 10-kernels dry weight (B) and increased dry weight (E) of developing kernels between WT and *zmbzip75-15.* Data were means± SD (*n* =4 biologically independent samples). (F) Measurement and comparison kernel moisture content between WT and *zmbzip75* mature kernels. Data are mean ± SD (*n* = 10 biologically independent samples). (G) and (H) Measurement and comparison of moisture content (G) and kernel dehydration rate (KDR) (H) of developing kernels between WT and *zmbzip75-15.* Data are mean ± SD (*n* =4 biologically independent samples). Statistical significance (*\*P < 0.05*; *\*\*P < 0.01*) was determined by two-tailed Student’s *t*-test.

Unexpectedly, the *zmbzip75* kernels appeared pale yellow and soft during the late stages of kernel development (from 30 to 39 DAP) compared to the wild-type (Figure 4A; Supplemental Figure S11), suggesting delayed dehydration and maturation. Indeed, the *zmbzip75* kernels had significantly higher kernel moisture content at harvest than the wild-type (Figure 4F). Dynamic measurement of kernel moisture content revealed that the *zmbzip75* kernels maintained higher moisture levels throughout kernel development, with particularly pronounced differences observed after 33 DAP (Figure 4G). Specifically, a markedly slower dehydration rate was observed in the *zmbzip75* kernels compared to the wild-type from 33 to 39 DAP (Figure 4H). These results collectively suggest that mutation of *ZmbZIP75* causes defects in grain filling and delayed grain dry-down.

### *zmbzip75* has less starch and lipid accumulation with developmental defects

The *zmbzip75* kernels could be easily distinguished from the wild-type on F2 ears by their smaller size and paler color starting at 10 DAP (Supplemental Figure S12). Light microscopy analysis of paraffin sections of 10 DAP and 15 DAP kernels revealed that the *zmbzip75* endosperm and embryo were significantly smaller than the wild-type, with the embryo being more severely affected (Figure 5A and 5B). The mutant endosperm exhibited markedly fewer starch granules and a narrower region of the under-expanded cells (UECs), which includes the aleurone and sub-aleurone layers (Figure 5C). The development of the BETL and the conducting zone (CZ) was also impacted in the *zmbzip75* kernels. At 10 DAP, the BETL of the mutant kernels had less cell wall ingrowths, and the cells in the CZ were more rounded, unlike the elongated cells observed in the wild-type (Figure 5D). Scanning electron microscopy analysis showed that the starch granules in the mature *zmbzip75* kernels were smaller (Figure 5E). Transmission electron microscopy further revealed that the *zmbzip75* endosperm cells contained fewer and smaller starch granules, while the number of protein bodies remained relatively unchanged (Figure 5F). Biochemical analyses revealed that the mutant kernels had significantly reduced starch and lipid contents but significantly increased soluble sugar content, compared to the wild-type. However, the total protein, zein, and nonzein contents did not show significant differences between the *zmbzip75* and wild-type kernels (Figure 5J; Supplemental Figure S13).

**Figure 5.**
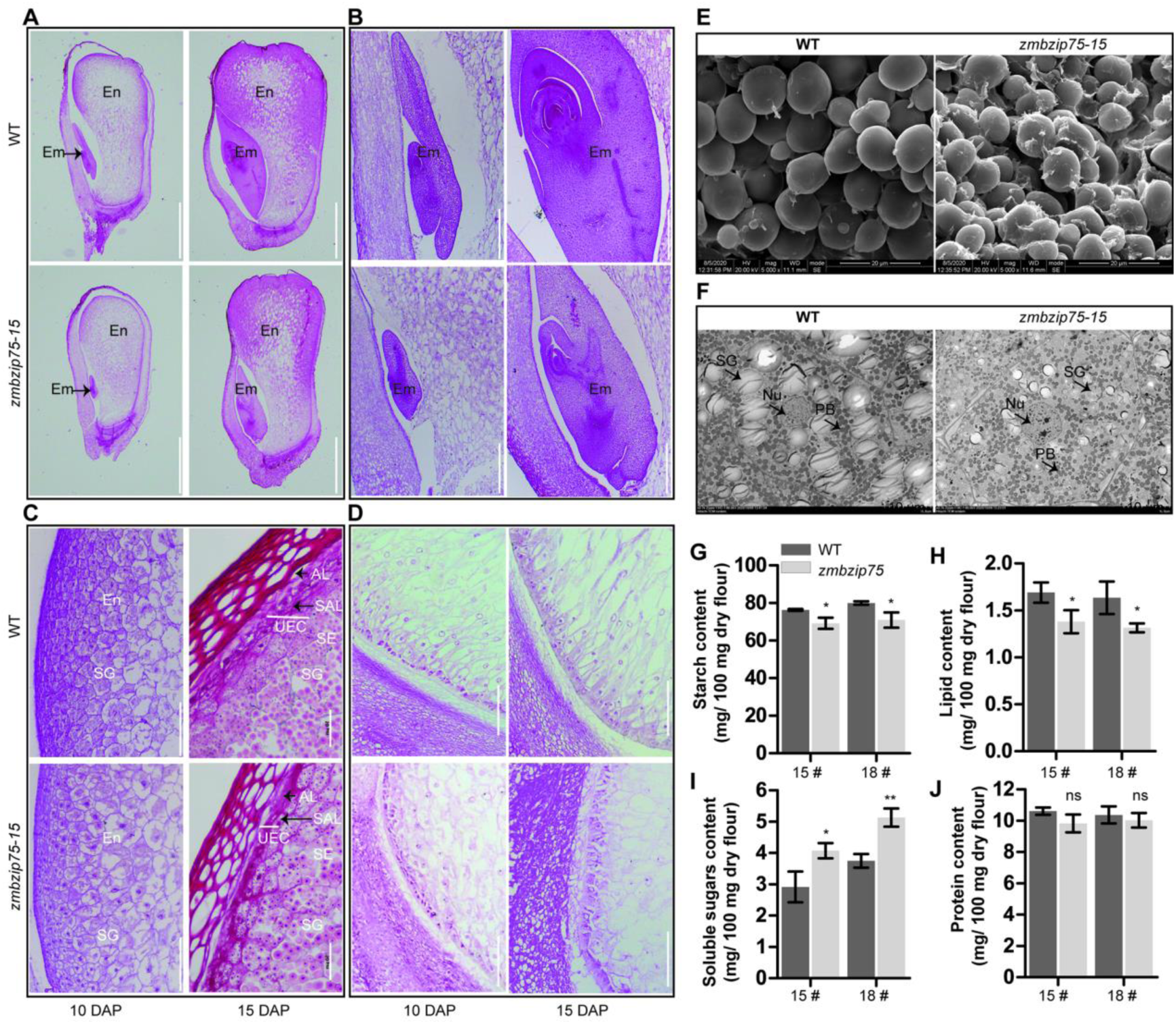
Cytological and biochemical analysis of wild-type and *zmbzip75* kernels. (A) to (D) Paraffin sections of developing kernels of WT and *zmbzip75-15* at 10-DAP and 15-DAP. En, endosperm.; Em, embryo; SG, starch granules; AL, aleurone; SAL, sub-aleurone; SE, starchy endosperm; UEC, under-expanded cells; CZ, conducting zone; PC, placento-chalazal region; PED, the vascular region of the pedicel; BETL, the basal endosperm transfer layer. A, Scale bar, 2 mm; B, Scale bar, 200 μm; C, Scale bar, 50 μm; D, Scale bar, 200 μm. (E) Scanning electron micrographs of the central regions of the mature endosperm of WT and *zmbzip75-15* kernel. Scale bar, 20 μm. (F) Transmission electron micrographs of starchy endosperm cells of 15-DAP WT and *zmbzip75-15* kernel. SG, starch granule; PB, protein body. Nu, nucleus. Scale bar, 10 μm. (G) to (J) Measurement and comparisons of starch content (G), lipid content (H), soluble sugars content (I) and protein content (J) of WT and *zmbzip75*. Error bars represent SD from three biological replicates. Statistical significance (*\*P < 0.05*; *\*\*P < 0.01*) was determined by two-tailed Student’s *t*-test.

To investigate the changes in metabolites of the *zmbzip75* kernels, we performed a nontargeted metabolomics analysis on 15 DAP kernels of the mutant and wild-type using LC-MS/MS. Principal component analysis showed that the six biological replicates of each sample clustered together under the positive and negative model (Supplemental Figure S14A). Statistical analysis identified 65 and 52 differentially accumulated metabolites in the positive and negative models, respectively (Supplemental Figure S14B; Supplemental Data Set S5). KEGG enrichment analysis of the significantly changed metabolites revealed that the “starch and sucrose metabolism” and “biosynthesis of unsaturated fatty acids” pathways were significantly enriched (Supplemental Figure S14C). Examining the starch synthesis pathway, we found that the levels of sucrose, glucose 6-phosphate (G6P), and glucose 1-phosphate (G1P) were significantly increased in the mutant kernels, suggesting impaired starch synthesis. In the fatty acid synthesis pathway, lysophosphatidic acid (LPA) and phosphatidic acid (PA) were significantly elevated, while phosphatidylcholine (PC) was significantly reduced (Supplemental Figure S14D and 14E). These results confirm that mutation of *ZmbZIP75* significantly affects both starch and lipid synthesis in the developing maize kernels.

### Transcriptomic alterations in the *zmbzip75* kernel

To decipher the molecular basis of ZmbZIP75 in kernel development and storage reserve synthesis, we performed transcriptome deep sequencing (RNA-Seq) analysis of the *zmbzip75* and wild-type kernels at 15 DAP (Supplemental Table S1). A total of 6,581 differentially expressed genes (DEGs) were identified (*P* value < 0.05, absolute fold change > 2.0), including 3,155 upregulated and 3,426 downregulated genes (Figure 6A; Supplemental Data Set S6). Gene ontology (GO) and Kyoto Encyclopedia of Genes and Genomes (KEGG) analyses revealed significant enrichment of DEGs in processes related to starch biosynthesis, starch and sucrose metabolism, sugar transmembrane transporter activity, fatty acid metabolism, response to water deprivation and DNA-binding TF activity (Figure 6B). Among the DEGs involved in starch and lipid synthesis, most of the endosperm-specific core SSRGs were significantly downregulated in the *zmbzip75* kernels (Figure 6C), consistent with observations in the ABA-deficient *dg1* mutant (Figure 1I). A number of lipid biosynthesis-related genes, including *acc1*, *acc2* and *oleosins*, were also notably downregulated (Figure 6C). *acc1* encodes an acetyl-CoA carboxylase, the first committed enzyme in the fatty acid synthesis pathway (Bates et al. 2013), while oleosins play a crucial role in maintaining oil body stability (Huang 1996). Additionally, several genes associated with aleurone and BETL development were significantly affected in the *zmbzip75* kernels (Figure 6C). For instance, *dek1*, encoding a membrane-localized protein essential for aleurone cell fate maintenance (Lid et al. 2002), and *mn1*, a BETL-specific cell wall invertase involved in sucrose cleavage and sugar transport to the kernels (Kang et al. 2009), were downregulated. Furthermore, the expression of key TFs related to kernel development and nutrient accumulation, such as *MRP1*, *MYBR29*, *NKD1/2*, *NAC128/130*, and *WRI1*, was also notably attenuated in the *zmbzip75* mutant (Figure 6C). The expression levels of these downregulated DEGs were validated using qRT-PCR in the wild-type and *zmbzip75* kernels, and the results were consistent with the RNA-Seq data (Supplemental Figure S15).

**Figure 6.**
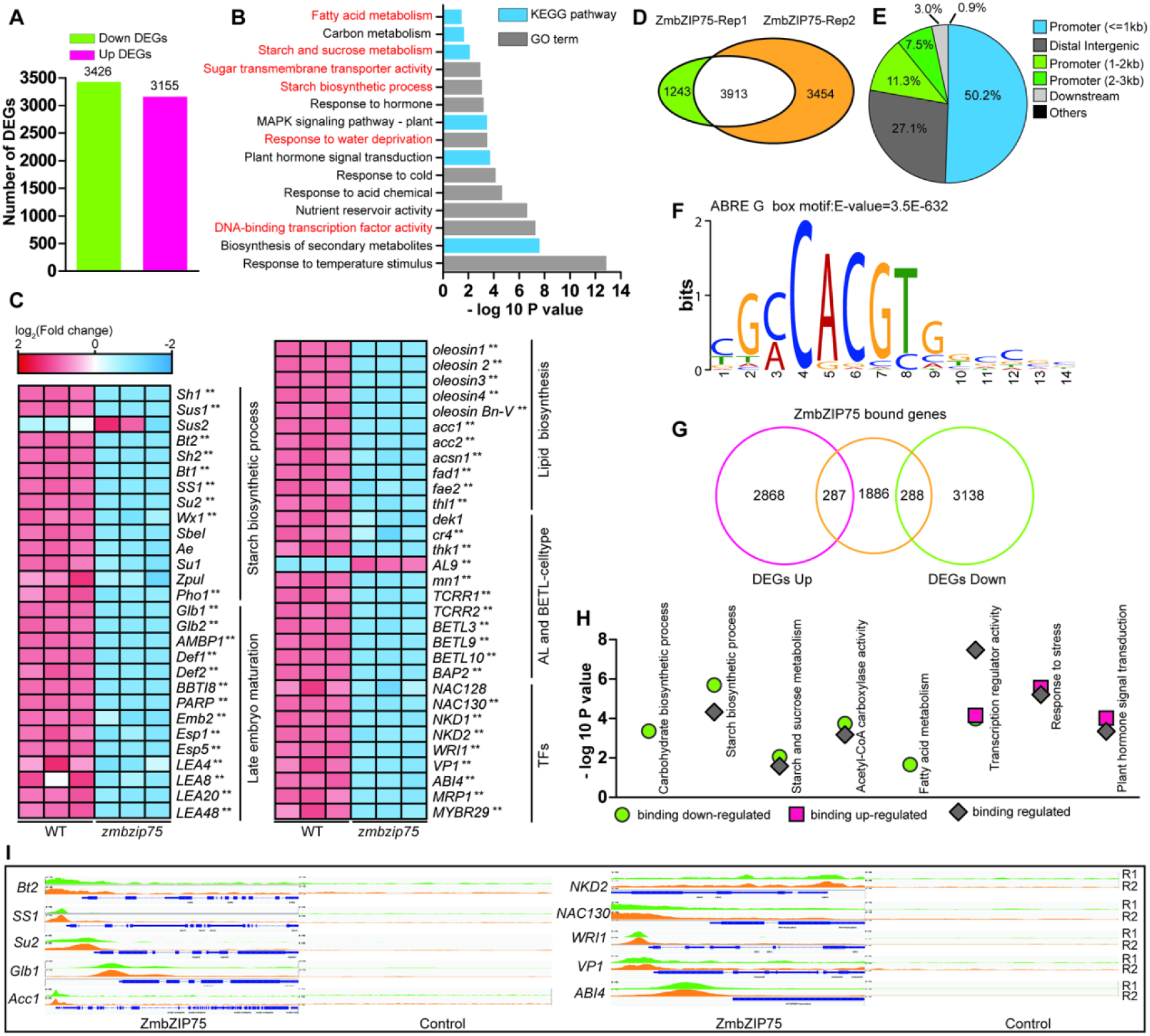
RNA-Seq analysis combined with ChIP-Seq analysis investigating the direct targets of ZmbZIP75. (A) Number of up-regulated and down-regulated genes in 15-DAP *zmbzip75-15* kernels compared to WT. (B) GO enrichment and KEGG enrichment analysis of DEGs. The GO term was shown as gray and KEGG pathway was shown as blue. (C) Heatmap showing the log_2_ (FKPM) of starch biosynthetic process related genes, highly embryo-expressed genes, lipid and fatty acid biosynthesis related genes, AL and BETL-celltype related genes, and some TFs related kernel development based on RNA-Seq. (D) Venn diagram showing the overlap of target genes identified from the two ChIP-Seq replicates. (E) Distribution of ZmbZIP75 binding regions in the maize genome. (F) The most significant elements of ZmbZIP75 according to MEME-ChIP analysis. The motif was identified by MEME-ChIP in 1.0 kb flanking sequences around the genic peaks, and enrichment P value was calculated using Fisher’s exact test. (G) Venn diagram showing the overlap of target genes identified from ChIP-Seq and RNA-Seq at 15-DAP. (H) Enrichment analysis of ZmbZIP75 directly regulated target genes in 15-DAP kernels. (I) Peak distributions of ZmbZIP75 binding sites for *Bt2*, *SS1*, *Su2*, *Glb1*, *Acc1*, *NKD2*, *NAC130*, *WRI1* and *VP1*. The loci were shown using the Integrated Genome Browser (IGV). Rep1 and Rep2 indicated the peak from the two ChIP-Seq replicates.

The downregulated DEGs also contained many embryo-specific genes related to late embryo maturation and grain desiccation, including *Glb1*, *Glb2*, *VP1*, *ABI4*, and *LEAs* (Figure 6C). VP1 is a central TF in grain desiccation, maturation and dormancy (Wilson et al. 1973; McCarty et al. 1991). Comparative analysis revealed that 684 genes were co-downregulated in the *zmbzip75* and *vp1* mutants (Zheng et al. 2019) (Supplemental Figure S16A), including numerous grain desiccation and maturation-related genes (Supplemental Figure S16B). Previous studies have shown that VP1 interacts with bZIP75 (Zhang et al. 2019a). We confirmed this interaction (Supplemental Figure S16C and 16D) and found that VP1 and bZIP75 synergistically induced the *Glb1* promoter activity (Supplemental Figure S16E-16G). These results indicate that bZIP75 modulates *VP1* transcription and interacts with VP1 to coordinately regulates grain desiccation and maturation in maize.

### ZmbZIP75 directly regulates genes involved in grain filling and maturation drying

To identify DEGs that are directly regulated by ZmbZIP75, chromatin immunoprecipitation sequencing (ChIP-Seq) analysis was performed with a bZIP75-specific antibody using 15 DAP wild-type kernels. Consistent peaks generated by two independent biological replicates were identified as binding sites. A total of 3913 common binding peaks were detected across the 10 chromosomes (Figure 6D; Supplemental Figure S17A; Supplemental Data Set S7), and predominantly located within the ≤1 kb (50.2%), 1-2 kb (11.3%), and 2-3 kb (7.5%) promoter regions upstream of TSS (Figure 6E). The binding sites were significantly concentrated near the TSS (Supplemental Figure S17B), consistent with the putative TF function of ZmbZIP75. A search for overrepresented motifs in the binding peaks within genic regions identified four G-box motifs (Supplemental Figure S17C), with the top-scoring motif (GCCACGTG) (E-value = 3.5E−632) matching a typical ABRE element (Figure 6F) (Hattori et al. 2002).

The binding peaks within the 3-kb promoter region corresponded to 2461 annotated genes, which were identified as putative direct targets of ZmbZIP75 (Supplemental Data Set S7). GO and KEGG analysis revealed that these targets were significantly enriched in processes related to starch biosynthesis, starch and sucrose metabolism, and response to water deprivation (Supplemental Figure S17D). Comparing of ZmbZIP75-bound targets with DEGs from the *zmbzip75* RNA-Seq data identified 575 direct targets, with 287 upregulated and 288 downregulated in the *zmbzip75* kernels (Figure 6G; Supplemental Data Set S8). GO and KEGG analysis of these direct targets highlighted significant enrichment in biological processes involved in starch and lipid synthesis, as well as transcription regulator activity (Figure 6H; Supplemental Table S2). Several endosperm-specific core SSRGs (*Bt2*, *SS1*, *Su2*, *Du1*, and *Wx*), lipid synthesis-related genes (*Acc1*, *Acc2*, *Fad1*, and *Oleosin*), and late embryo maturation-related genes (*Glb1*, *AMBP1*, *Def2*, *BBTI8*, and *LEAs*) were identified as direct targets of ZmbZIP75 and were significantly downregulated in the *zmbzip75* kernels. Additionally, ZmbZIP75 was found to directly regulate multiple known key TFs essential for kernel development and grain filling, such as *NKD2*, *NAC130*, *WRI1*, *ABI4*, and *VP1* (Figure 6I).

We further validated the binding and transactivation of ZmbZIP75 to these target genes. An ABRE element was identified at positions -152 bp and -388 bp upstream of the translation start site (ATG) in the *Bt2* and *Su2* promoters, respectively. ChIP-qPCR analysis using 15 DAP kernels with a ZmbZIP75-specific antibody revealed that the ABRE-containing fragments in the *Bt2* and *Su2* promoters were notably enriched in the wild-type compared to the *zmbzip75* mutant (Figure 7A). Electrophoretic mobility shift assay (EMSA) confirmed that ZmbZIP75 specifically bound to the ABREs in the *Bt2* and *Su2* promoters (Figure 7B). We then performed transient expression assays to determine whether ZmbZIP75 directly activated the *Bt2* and *Su2* promoter activities. A reporter plasmid containing a *LUC* gene driven by either *Bt2* or *Su2* promoter was co-transformed with an effector plasmid expressing *ZmbZIP75* driven by the *ubiquitin* (*ubi*) promoter. The *ubi* promoter driving β-glucuronidase (GUS) was used as an internal control. Co-transformation of the reporter and effector plasmids into developing endosperm significantly enhanced the *Bt2* and *Su2* promoter activities, compared to the reporter plasmid alone (Figure 7C), suggesting that ZmbZIP75 directly activated *Bt2* and *Su2* transcription.

**Figure 7.**
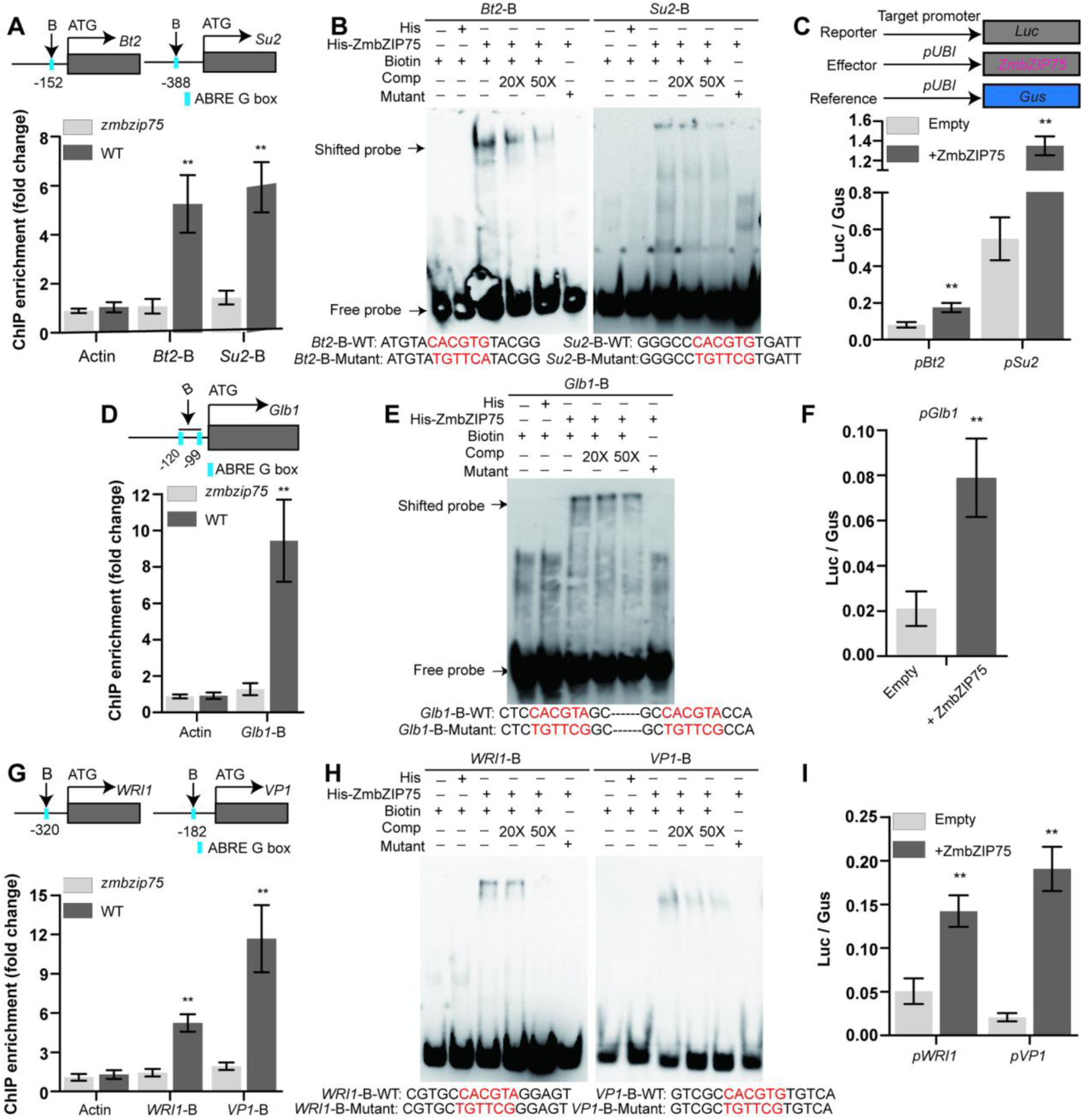
ZmbZIP75 directly binds to the promoters of target genes for transactivation. (A), (D) and (G) ChIP-qPCR assay showing the binding of ZmbZIP75 to the promoter of *Bt2* and *Su2* (A), *Glb1* (D) and *WRI1* and *VP1* (G) in vivo. The examined regions were marked in blue under the schematic diagrams of the target gene loci. Error bars represent SD from three biological replicates. (B), (E) and (H) EMSA showing the direct binding of ZmbZIP75-His to the promoters of *Bt2* and *Su2* (B), *Glb1* (E) and *WRI1* and *Glb1* (H). The individual wild-type probe was biotin-labelled and incubated with the recombinant protein ZmbZIP75-His. The His protein alone was used as negative controls. The mutant motifs are indicated in red. Comp, competing probes unlabeled with biotin. (C), (F) and (I) The transactivation of *pBt2* and *pSu2* in maize endosperms (C), the activity of *pGlb1* (F) and the activity of *pWRI1* and *pVP1* (I) in maize embryos by ZmbZIP75 via particle bombardment. Error bars represent SD from three biological replicates. Statistical significance (*\*P < 0.05*; *\*\*P < 0.01*) was determined by two-tailed student’s *t*-test.

*Glb1* encodes a globulin protein that accumulates in the embryo during maturation drying stage (Belanger and Kriz 1989). Globulin accumulation depends on the presence of ABA and has served as a marker for the study of late embryo development and seed maturation in maize (Kriz et al. 1990; Rivin and Grudt 1991). Two ABREs were found to be located at positions -120 bp and -99 bp upstream of the ATG. ChIP-qPCR and EMSA assays revealed that ZmbZIP75 specifically bound to these two ABREs both *in vivo* and *in vitro* (Figure 7D and 7E). Transient expression assays further showed a significant enhancement of *Glb1* promoter activity by ZmbZIP75 (Figure 7F).

A group of TFs, including *NKD2*, *NAC130*, *WRI1*, and *VP1*, were also identified as direct targets of ZmbZIP75 (Supplemental Table S2). Notably, NKD2 (with its paralog NKD1) was recently reported to participate in controlling the transition from endosperm cell differentiation to grain filling (Wu et al. 2023), while NAC130, together with its paralog NAC128, was shown to coordinate the nutrient uptake in the BETL and storage reserve synthesis in the endosperm to facilitate grain filling (Chen et al. 2023). Furthermore, WRI1 has been identified as a master regulator involved in the regulation of oil biosynthesis in the maize embryo (Pouvreau et al. 2011). Through ChIP-qPCR, EMSA and transient expression analyses, ZmbZIP75 was found to specifically bind to the respective ABRE present in the *WRI1* and *VP1* promoters and enhance their promoter activities (Figure 7G-7I). Similarly, ZmbZIP75 recognized the ABREs in the *NKD2* and *NAC130* promoters and transactivated their expression, as revealed by ChIP-qPCR and transient expression analyses (Supplemental Figure S18A and 18B).

In summary, these finding demonstrate that ZmbZIP75 directly transactivates a series of grain filling and dehydration-related genes, as well as key TFs, indicating it likely functions as a central TF in regulating maize kernel development.

### ZmSnRK2.10 phosphorylates and stabilizes ZmbZIP75 in the embryo

ZmbZIP75 primarily exists in a phosphorylated form during late embryo development (Figure 3G). To identify kinases that interact with ZmbZIP75, total proteins were extracted from 15 DAP maize B73 kernels and immunoprecipitated using a ZmbZIP75-specific antibody. The precipitated proteins were then analyzed by liquid chromatography–tandem mass spectrometry (LC-MS/MS), which identified a candidate protein with high coverage belonging to the subclass III SnRK2 family, ZmSnRK2.10 (Supplemental Figure S19A). Subclass III SnRK2s are core components of the ABA signaling pathway, phosphorylating downstream TFs to activate ABA-responsive gene expression (Kobayashi et al. 2005). Our previous study showed that *ZmSnRK2.10* is primarily expressed in the middle and late stages of kernel development (Long et al. 2021), consistent with the expression pattern of *ZmbZIP75* (Figure 3A). To verify the direct interaction between ZmbZIP75 and ZmSnRK2.10, yeast two-hybrid assays were performed. Rapid growth of the yeast strain harboring bait BD-SnRK2.10 and prey AD-bZIP75 was observed on selective media, compared to the negative controls (Figure 8A). Further split-LUC complementation and GST pull-down assays confirmed that ZmSnRK2.10 directly interacts with ZmbZIP75 *in planta* (Figure 8B and 8C).

**Figure 8.**
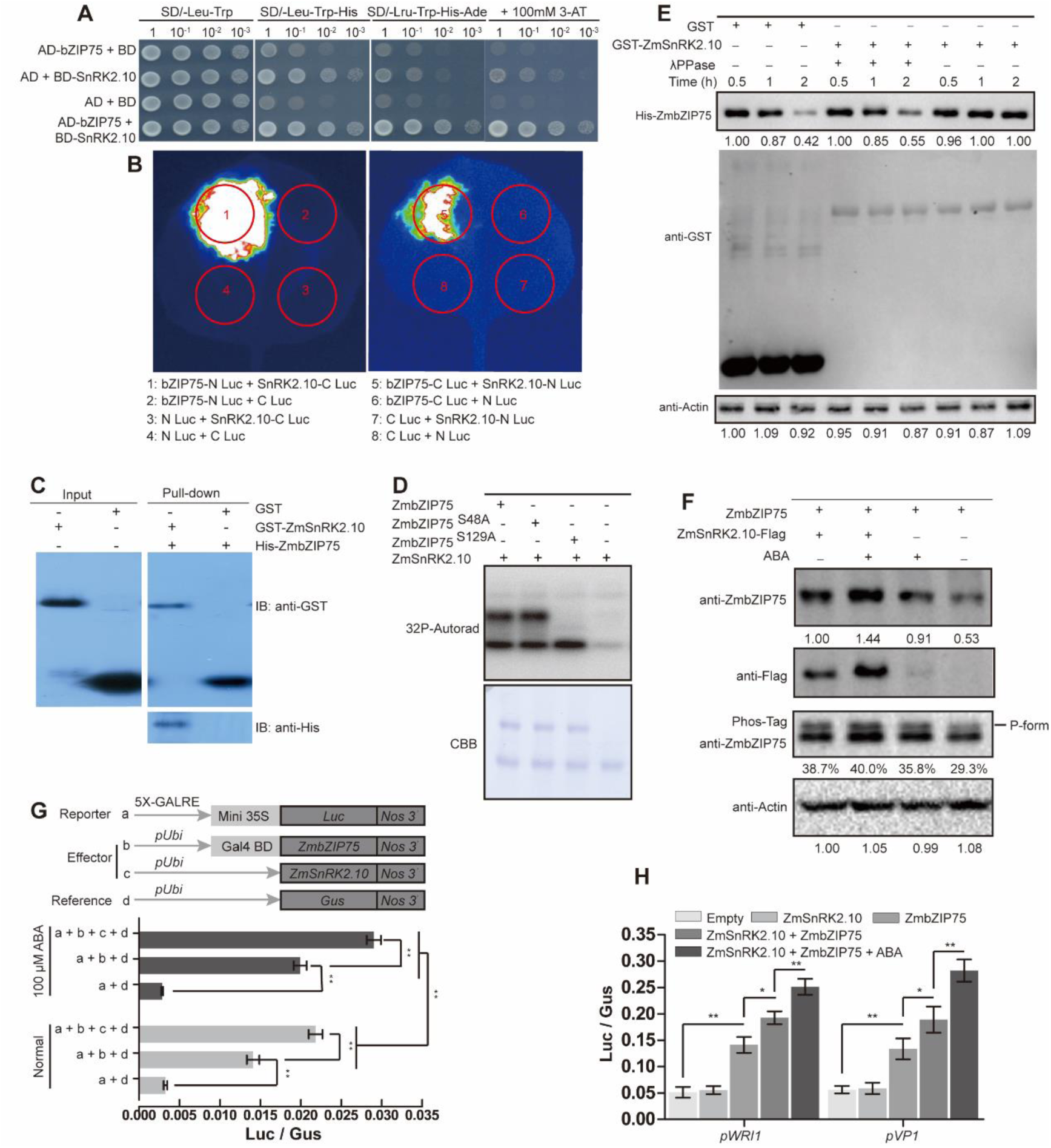
ZmSnRK2.10 interacts with and phosphorylates ZmbZIP75 to enhance its transactivation ability. (A) The interaction between ZmSnRK2.10 and ZmbZIP75 in yeast. Transformed yeast cells (1,1/10,1/100,1/1000 cellular dilutions) were grown on SD/-Leu-Trp medium, selected on SD/-Leu-Trp-His, SD/-Leu-Trp-His-Ade, and SD/-Leu-Trp-His-Ade + 100 mM 3AT inhibitor. (B) The interaction between ZmSnRK2.10 and ZmbZIP75, as tested by Split-Luciferase in *N. benthamiana*. The LUC intensity was measured through co-expression of different combinations. (C) In vitro pull-down assay showing the interaction between ZmSnRK2.10 and ZmbZIP75. Recombinant GST and GST-ZmSnRK2.10 were immobilized on glutathione agarose beads and incubated with His-ZmbZIP75 protein. The immunoblotting was detected by anti-His and anti-GST antibodies. (D) ZmbZIP75 is phosphorylated by ZmSnRK2.10 at Ser129. GST-ZmSnRK2.10, His-ZmbZIP75^S48A^ and His-ZmbZIP75^S129A^ were expressed in *E.coli* and purified, then subjected to an in vitro phosphorylation assay (top), the recombinant proteins were detected by Coomassie brilliant blue (CBB) staining (bottom). (E) ZmSnRK2.10 stabilizes ZmbZIP75 protein in cell-free degradation assays. Total proteins extracted from WT kernels at 18 DAP were incubated with equal amounts of His-ZmbZIP75 with GST-ZmSnRK2.10 or GST for 0.5h, 1h, and 2h. λPPase treatment was applied to dephosphorylate ZmbZIP75. Actin was used as loading control. ImageJ software was used to quantify the immunoblot results. (F) ABA enhances the phosphorylation and stability of ZmbZIP75 protein in maize embryos. The embryos at 18 DAP were transformed with or without *ZmSnRK2.10-Flag* plasmid DNA by particle bombardment, then under the absence or presence of ABA (100 μM). Extracting an equal amount of total protein, the phosphorylation and stability of ZmbZIP75 protein were determined by Phos-Tag and immunoblot, respectively. Actin was used as a loading control. (G) ZmSnRK2.10 mediates ABA to enhance the transactivation activity of ZmbZIP75. Error bars represent SD from three biological replicates. (H) Luc/Gus assay showing that ZmSnRK2.10 and ABA enhance transactivation capacity of ZmbZIP75 on the *ZmWRI1* and *VP1* in embryos. Error bars represent SD from three biological replicates. Statistical significance (*\*P < 0.05*; *\*\*P < 0.01*) was determined by two-tailed student’s *t*-test.

*In vitro* kinase assays using recombinant GST-ZmSnRK2.10 and His-ZmbZIP75 proteins revealed that ZmbZIP75 is phosphorylated by ZmSnRK2.10 (Supplemental Figure S19B). ABA-activated SnRK2s are known to phosphorylate Ser/Thr residue within R-X-X-S/T motifs (Furihata et al. 2006). Two such motifs with putative phosphorylation sites at Ser48 and Ser129 were identified in the conserved domains of ZmbZIP75 (Supplemental Figure S9A). To test whether these two sites can be phosphorylated by ZmSnRK2.10, we introduced two point mutations in ZmbZIP75 that replaced Ser48 and Ser129 with the nonphosphorylatable alanines (Ala), respectively (named ZmbZIP75^S48A^ and ZmbZIP75^S129A^). *in vitro* phosphorylation assays showed that no phosphorylation signal was detected when Ser129 was mutated to Ala, while ZmbZIP75^S48A^ did not affect phosphorylation, indicating that ZmSnRK2.10 specifically phosphorylates ZmbZIP75 at Ser129 (Figure 8D). In addition, ZmSnRK2.10 kinase activity was reported to be activated by ABA (Long et al. 2021). To investigate the effect of ABA on ZmbZIP75 phosphorylation, Phos-tag SDS-PAGE assay were performed. The results showed that ABA treatment of 18 DAP maize kernels significant enhanced both the protein abundance and phosphorylation level of ZmbZIP75 (Supplemental Figure S19C).

To elucidate the biological significance of ZmbZIP75 phosphorylation by ZmSnRK2.10, we first performed an *in vitro* cell-free degradation assay to test whether ZmSnRK2.10-mediated phosphorylation affects the protein stability of ZmbZIP75. When recombinant His-ZmbZIP75 was incubated with total proteins from 18-DAP wild-type kernels, the protein level of His-ZmbZIP75 was significantly decreased after 2 h. Co-incubation with GST-ZmSnRK2.10 significantly attenuated the degradation of His-ZmbZIP75. However, ZmbZIP75 degradation still occurred upon the addition of λPPase, even in the presence of GST-ZmSnRK2.10 (Figure 8E). To investigate the impact of ABA and ZmSnRK2.10 on the protein stability of ZmbZIP75 *in vivo*, a Flag-tagged ZmSnRK2.10 construct was transiently transformed into developing maize embryos by particle bombardment. The bombarded embryos were treated with or without ABA and then subjected to immunoblot and Phos-tag analyses. The results showed that the stability of ZmbZIP75 was significantly enhanced in both the ZmSnRK2.10-bombarded and the ABA-treated embryos. Notably, the combination of ZmSnRK2.10 transformation and ABA treatment led to a further increase in ZmbZIP75 stability (Figure 8F). These results collectively demonstrate that ABA-activated ZmSnRK2.10 phosphorylates ZmbZIP75 and thereby enhances its protein stability.

Given that the N-terminal region (1-162 amino acids) containing Ser129 is essential for the transactivation activity of ZmbZIP75 (Figure 3F), the effect of ZmSnRK2.10 on ZmbZIP75 transactivation was investigated by a dual-luciferase reporter system. Co-transformation of ZmbZIP75 with ZmSnRK2.10 in developing embryos significantly enhanced the transactivation activity of ZmbZIP75, and this enhancement was further amplified by ABA treatment (Figure 8G). However, the substitution of Ser129 to Ala in ZmbZIP75 abolished the enhancement effect by ZmSnRK2.10 (Supplemental Figure S20A). Transient expression assays also showed that co-transformation of ZmbZIP75 and ZmSnRK2.10 in developing embryos, combined with ABA treatment, jointly enhanced the activities of the *WRI1* and *VP1* promoters (Figure 8H).

Taken together, these results highlight the importance of the SnRK2.10-ZmbZIP75 module in mediating ABA signaling to regulate late grain maturation in maize.

### Overexpression of *ZmbZIP75* enhances maize yield and accelerates grain dry-down

Mutant analysis showed that ZmbZIP75 plays an important role in grain filling and late maturation processes (Figure 4). To explore the potential of ZmbZIP75 for breeding applications, three overexpression lines driven by the *ubi* promoter were generated (Supplemental Figure S21A). qRT-PCR and immunoblot analyses showed a significant increase in ZmbZIP75 transcripts and protein levels in these overexpression lines (Supplemental Figure S21B and 21C). Comparing the grain development dynamics between wild-type and overexpression lines revealed that from the onset of grain filling (12 DAP) to maturity, the kernel size of the overexpression lines was significantly larger than that of the wild-type (Supplemental Figure S22A). At 15 DAP, the kernel width and thickness of *ZmbZIP75* overexpression lines were notably increased, with endosperm cells appearing to accumulate more starch granules (Supplemental Figure S21G and 21H). Consequently, the *ZmbZIP75*-overexpressed kernels at 15 DAP exhibited higher fresh weight and starch content (Supplemental Figure S21I and 21J).

Phenotypic analysis of mature kernels revealed that the *ZmbZIP75*-overexpressed kernels had significantly increased kernel width and thickness, while kernel length remained unchanged compared to the wild-type (Figure 9A). The 100-kernel weight and starch content were also significantly higher in the overexpression lines (Figure 9B and 9C). The lipid content showed a slight but not statistically significant increase (Figure 9D). However, due to the substantial increase in grain weight, the lipid content per grain was significantly higher in the overexpression lines compared to the wild-type. Analysis of grain dry weight accumulation dynamics revealed a significantly higher grain filling rate throughout the effective grain filling period (12-33 DAP) in *ZmbZIP75* overexpression lines compared to wild-type (Figure 9E; Supplemental Figure S22B).

**Figure 9.**
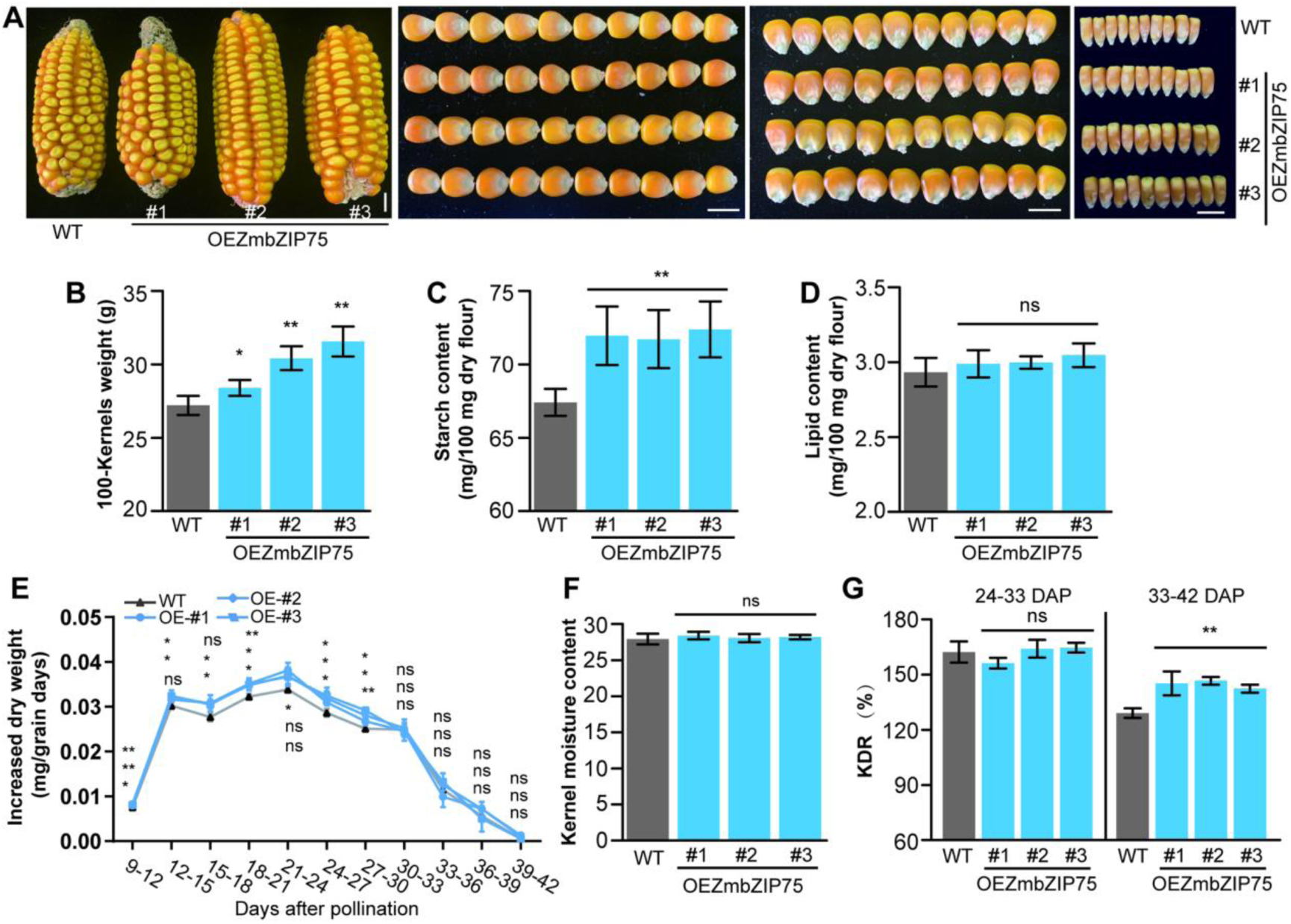
Phenotypes and biochemical analysis of *ZmbZIP75*-overexpression kernels. (A) The ear phenotype and kernel size of the wild-type B104 (WT) and *ZmbZIP75*-overexpression lines (#1, #2, #3). Scale bar, 1 cm. (B) to (D) Measurement and comparisons of 100-kernels weight (B), starch content (C) and lipid content (D) of WT and OE kernels. (E) Measurement and comparisons of the increased dry weight of WT and OE developing kernels. (F) Measurement and comparisons of the kernel moisture content of WT and OE kernels. (G) Quantification of KDR between the WT and OE kernels from 24 DAP to 42 DAP. Data were means± SD. Statistical significance (*\*P < 0.05*; *\*\*P < 0.01*) was determined by two-tailed Student’s *t*-test.

Previous studies have indicated that large-grain maize varieties typically have higher kernel moisture content (Barnwal et al. 2012; Li et al. 2021b). Surprisingly, the moisture content of *ZmbZIP75-*overexpressed kernels did not differ significantly from that of the wild-type (Figure 9F). Further analysis of the kernel moisture content dynamics from 24-36 DAP showed significantly higher moisture content in *ZmbZIP75*-overexpressed kernels compared to the wild-type. However, this increase gradually diminished during the maturation drying stage (33-42 DAP), with no difference observed at harvest (42 DAP) (Supplemental Figure S22C). Examination of the kernel dehydration rates 6 days before and after 33 DAP revealed no significant difference in dehydration rate during 24-33 DAP between *ZmbZIP75-*overexpressed and wild-type kernels. However, during 33-42 DAP, the dehydration rate of *ZmbZIP75-*overexpressed kernels was significantly higher than that of the wild-type (Figure 9G). Additionally, *ZmbZIP75* overexpression did not alter plant architecture, with no significant differences in plant height or ear height (Supplemental Figure S21D-21F). These findings suggest that overexpression of *ZmbZIP75* significantly increases the grain filling rate and the kernel dehydration rate during maturation drying, resulting in higher grain yield without increasing kernel moisture content.

To evaluate the value of *ZmbZIP75* overexpression in hybrids, eight hybrids were generated by crossing wild-type and overexpression line OE-3 (the highest *ZmbZIP75* expression level among three OE lines) with four elite inbred lines: Zheng58, Chang7-2, PH6WC, and PH4CV. Zheng58 and Chang7-2 are parents of the Zhengdan958, a traditional high-yielding hybrid widely planted in China but known for high grain moisture content at harvest. PH6WC and PH4CV are parents of Xianyu335, a transitional high-yielding hybrid with relatively lower grain moisture content (Wang et al. 2019a). As expected, all *ZmbZIP75* overexpression hybrids showed significantly larger kernel size compared to the corresponding wild-type hybrids, with increased kernel width and thickness, but no significant change in kernel length, consistent with the results from the overexpression lines (Figure 10A-10D; Supplemental Figure S23I-23K). Except for the OE/B104 × Zheng58 hybrid, which showed a slight but not statistically significant increase in 100-kernel weight compared to the wild-type B104 × Zheng58 hybrid, the other three *ZmbZIP75* overexpression hybrids (OE/B104 × Chang7-2, OE/B104 × PH6WC, and OE/B104 × PH4CV) had significantly higher 100-kernel weight than their respective wild-type hybrids, with increases of 11%, 4.6%, and 6.7%, respectively (Figure 10E).

**Figure 10.**
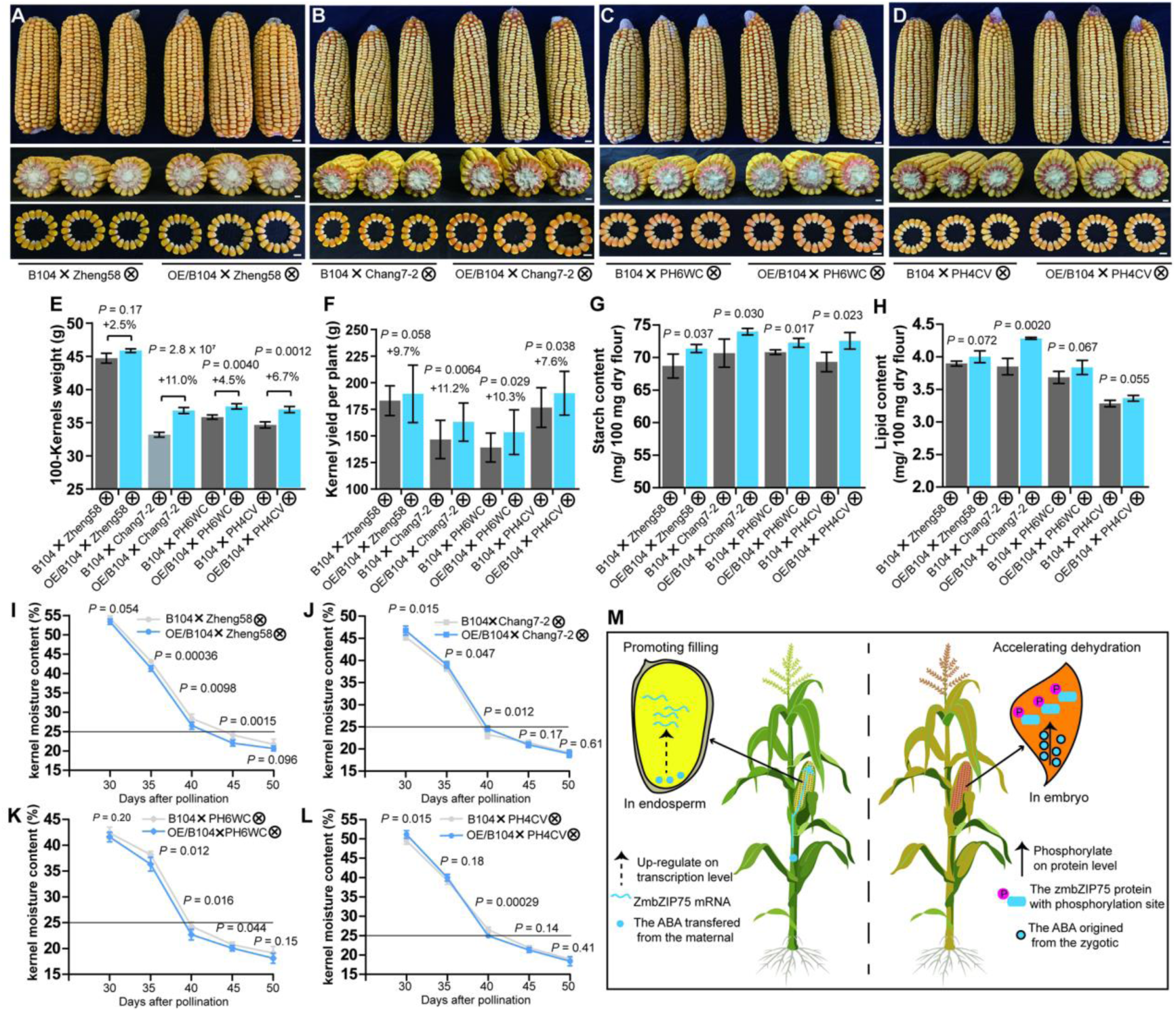
ZmbZIP75 coordinates grain filling and kernel dehydration in maize. (A) to (D) Ear and mature kernel phenotypes of F_1_ hybrids derived from overexpression line OE3 and the background B104 by crossing to four representative inbred lines (F_1_ hybrids derived from overexpression line OE3 and the corresponding no transgenic (NT) control by crossing to four elite inbred lines (Zheng58 (A), Chang7-2 (B), PH6WC (C), PH4CV (D)). (E) to (H) Measurement and comparisons of the 100-kernels weight (E, *n* = 16-20), kernel yield per plant (F, *n* = 16-20), starch content (G, *n* = 3) and lipid content (H, *n* = 3) of four F_1_ hybrids. (I) to (L) Measurement and comparisons of the moisture content of four F_1_ hybrids during maturation (*n* = 6). Statistical significance (*\*P < 0.05*; *\*\*P < 0.01*) was determined by two-tailed student’s *t*-test. (M) Proposed model of ZmbZIP75 regulating the maize seed development. In endosperm filling stage, ZmbZIP75 expression is up-regulated by ABA, which primarily originates from the maternal plant. ZmbZIP75 directly binds to the ABRE G-box in the promoter of *Bt2*, *SS1*, *Su2, NKD2* and *NAC130*, etc., to enhance their expressions, leading to promote endosperm development and grain filling. During seed embryonic and maturation, when ABA contents are substantially increased from zygotic origin, ABA-activated ZmSnRK2.10 interacts with and phosphorylates ZmbZIP75 at Ser129 to promote its stability and transactivation capacity in embryo. ZmbZIP75 directly binds to the ABRE G-box in the promoter of *LEAs*, *WRI1*, *VP1* and *ABI4*, etc., to enhance their expressions, leading to promote embryo development and dehydration maturation.

The grain yield per plant was increased by 9.7%, 11.2%, 10.3%, and 7.6% in all four *ZmbZIP75* overexpression hybrids, respectively, compared to the wild-type hybrids (Figure 10F). Importantly, no significant changes were observed in other agronomic traits such as plant architecture, plant height, ear height, ear length, and ear width (Supplemental Figure S23A-23H), suggesting that *ZmbZIP75* overexpression may increase maize yield per unit area under similar cultivation conditions. Additionally, the kernel starch content was significantly increased in all *ZmbZIP75* overexpression hybrids (Figure 10G). The kernel lipid content was also substantially increased, although only the OE/B104 × Chang7-2 hybrid showed a statistically significant difference (Figure 10H).

We investigated the dynamic changes in kernel moisture content during the maturation drying stage (30-50 DAP) between *ZmbZIP75* overexpression hybrids and their corresponding wild-type hybrids. It was observed that OE/B104 × Zheng58 and OE/B104 × PH6WC maintained lower kernel moisture content throughout the entire maturation drying stage compared to the wild-type hybrids (Figure 10I and 10K). The OE/B104 × PH4CV hybrid also exhibited lower kernel moisture around 40 DAP compared to the wild-type (Figure 10L). In contrast, the OE/B104 × chang7-2 hybrid had significantly higher kernel moisture content between 30-40 DAP than the wild-type (Figure 10L), possibly due to its dramatically increased kernel size and weight (Figure 10E). Based on the standard that a 25% kernel moisture content at harvest can ensure the quality of maize mechanical grain harvesting (Plett 1994; Gao et al. 2021), we found that the OE/B104 × Zheng58, OE/B104 × PH6WC, and OE/B104 × PH4CV hybrids required less time to reach this threshold compared to their wild-type hybrids (Figure 10I, 10K and 10L).

In summary, these results demonstrate that *ZmbZIP75* overexpression significantly increases grain yield and reduces kernel moisture content in maize hybrids. This finding could facilitate the development of new maize varieties that are both high-yielding and suitable for mechanical grain harvesting.

## Discussion

### Conserved and divergent functions of ABI5 homologs in plant seed development

The phytohormone ABA regulates multiple aspects of plant seed development, including endosperm differentiation, storage compound accumulation, and seed desiccation and dormancy (Ali et al. 2022). ABI5, a member of the A-group bZIP TF family, is phosphorylated by SnRK2s and serves as a crucial component of ABA signaling cascades, mediating ABA responses by binding to the ABRE elements (Finkelstein and Lynch 2000; Nakashima et al. 2009). In Arabidopsis, ABI5 controls embryo cell number and seed size by repressing the expression of *SHB1*, a key gene involved in early seed development. The *abi5* mutant exhibits significantly increased seed size and weight (Cheng et al. 2014). During late seed development, phosphorylated ABI5 activates the expression of late embryo maturation-related genes, such as LEAs, indicating its role in seed desiccation tolerance (Nakashima et al. 2009). However, the *abi5* mutant did not show obvious seed dormancy phenotypes in Arabidopsis (Finkelstein 1994). In contrast, mutation of *ABI5* in legumes like *Medicago truncatula* and pea (*Pisum sativum*) significantly affects seed dormancy and longevity, suggesting species-specific roles of ABI5 in regulating seed dormancy. Notably, *abi5* mutants in these species do not affect seed size or weight, implying a scarce role for ABI5 in seed filling (Zinsmeister et al. 2016).

ZmbZIP75, a homolog of Arabidopsis ABI5 (Supplemental Figure S6A), is predominantly expressed in maize kernels, with the highest expression during late embryo development (Figure 3A-3C and 3G), similar to ABI5 in Arabidopsis (Finkelstein and Lynch 2000). Unlike its dicot homologs, which are not involved in seed filling, ZmbZIP75 expression is closely associated with maize HGW (Figure 2). Knockout of *ZmbZIP75* significantly reduces starch accumulation and grain filling rate, leading to decreased grain size and weight (Figure 4A-4E). The reasons for the distinct roles of ABI5 homologs in seed filling between maize and legumes are not yet clear. At least it is unlikely due to differences in seed starch content, as ABI5 does not regulate seed filling in starch-rich pea (Zinsmeister et al. 2016).

Mutation of *ZmbZIP75* also significantly reduces the kernel dehydration rate during late kernel development, resulting in higher kernel moisture content at harvest compared to the wild-type (Figure 4F-4H). ZmbZIP75 directly regulates VP1 and interacts with it to co-regulate the expression of a series of maturation-related genes (Figure 7G-7I; Supplemental Figure S16). Similarly, mature kernels of the *vp1* mutant also exhibit significantly increased moisture content (Neill et al. 1986), indicating that ZmbZIP75 and VP1 synergistically control the kernel dehydration process. However, the specific roles of the genes co-regulated by bZIP75 and VP1 in kernel dehydration require further investigation. ZmbZIP75 also regulates the expression of dehydrated protectant LEAs (Figure 6C), similar to ABI5 in Arabidopsis and legumes, suggesting a conserved function of ABI5 homologs in seed desiccation tolerance. However, the seed dehydration rate and moisture content of *abi5* mutants in legumes do not differ from the wild-type (Zinsmeister et al. 2016), indicating that legume ABI5 may require additional regulatory factors to cooperatively control seed dehydration. Alternatively, the seed dehydration process in legumes may be governed by fundamentally distinct regulatory pathways compared to that in maize.

### ZmbZIP75 is a core regulator of grain filling and kernel dehydration through integration of maternal-derived and zygotic ABA signaling

ABA accumulation exhibits two distinct peaks during maize kernel development. One peak occurs in the lag phase, and another appears just prior to physiological maturity (PM). However, ABA remains at a basal level throughout the grain filling period (Jones and Brenner 1987). The ABA accumulated in developing endosperm during the lag phase and grain filling period is derived from the maternal plant. In contrast, ABA accumulated in the late stage of kernel development is synthesized *in situ* within the kernel (Ober and Setter 1992). This distinction is further supported by observations that mutation of *DG1*, which blocks ABA transport from maternal tissues to kernels, results in significantly decreased kernel ABA content during the grain filling period (18 DAP), while ABA level at late maturation (36 DAP) is similar to the wild-type (Supplemental Figure S2B). The decreased grain ABA content in the *dg1* mutant impairs starch synthesis and ultimately affects kernel weight, indicating that the basal-level maternal ABA is crucial for maintaining normal maize grain filling (Figure 1F-1I). The basal level of ABA signaling is essential for normal plant growth and development under non-stressed conditions (Yoshida et al. 2019). For instance, mutations in group I ABA receptors (*PYL1/6/12*) have been showed to improve plant growth and grain yield in rice (Miao et al. 2018).

Under moderate soil drying or ABA treatment, the appropriately increased ABA in grains significantly promotes starch synthesis and grain filling in crops (Yang et al. 2001; Yang et al. 2004; Zhang et al. 2012; Yuan et al. 2023; Yu et al. 2024). In contrast, exogenous ABA supply at high concentrations results in decreased grain filling rate and grain weight (Zhu et al. 2011; Zhang et al. 2017a). The present study also reveals that grain weight and starch content in developing maize kernels were significantly enhanced by moderate ABA treatment (100 μM), while no obvious enhancement effect was observed under high concentration ABA treatment (300 μM) (Figure 1A-1C), indicating that a moderate increase in basal-level ABA in developing kernels can promote grain filling and final grain yield.

*ZmbZIP75* expression was significantly induced under moderate ABA treatment (100 μM) but not under high concentration (300 μM) (Supplemental Figure S4A and 4B), similar to the trend observed in grain weight and starch content (Figure 1A-1C). ZmbZIP75 accumulates in the non-phosphorylated state in developing endosperm (Figure 3G), and directly transactivates several core SSRGs and key TFs, such as *NKD2* and *NAC130*, to regulate endosperm starch synthesis (Figure 6I, Figure 7A-7C; Supplemental Figure S18; Supplemental Table S2). ZmbZIP75 also indirectly activates BETL-specific genes and two key TFs, *MRP1* and *MYBR29* (Figure 6C), both of which are essential for BETL differentiation and development (Gomez et al. 2009; Yuan et al. 2024), suggesting that ZmbZIP75 may indirectly regulate assimilate transport to the kernels. Unlike known endosperm regulators such as *O2*, *PBF*, and *bZIP22*, which mainly control zein synthesis (Zhang et al. 2015; Li et al. 2018a), and *NAC128/130*, which directly regulate both starch and zein synthesis (Chen et al. 2023), ZmbZIP75 primarily controls starch synthesis in the endosperm without significantly affecting protein accumulation (Figure 5E-5G and 5J; Supplemental Figure S13).

Recent studies have revealed that two TFs ZmABI19 and ZmbZIP29, which are predominantly expressed during early kernel development, are phosphorylated by SnRK2.2, facilitating early ABA signaling to initiate grain filling by transactivating several key TFs such as *O2*, *PBF*, *O11*, *bZIP22*, and *NAC130* (Yang et al. 2021; Yang et al. 2022). Following the initiation of grain filling, rapid starch accumulation requires sustained high-level expression of a set of endosperm-specific SSRGs throughout the grain filling period (Hu et al. 2021). Our present study demonstrates that ZmbZIP75 functions as a core regulator mediating basal ABA signaling in developing endosperm, maintaining high expression of these SSRGs, and thereby promoting starch synthesis and grain filling. The regulatory mechanisms underlying ABA-inducible *ZmbZIP75* transcription remain unclear. Association analysis showed that the elite haplotype Hap.B (SNP250-C) is highly correlated with HGW and *ZmbZIP75* expression (Figure 2D and 2F). This SNP is located within a putative ABRE element in the *ZmbZIP75* promoter (Figure 2C), indicating that *ZmbZIP75* transcription may be auto-activated or activated by other A-group bZIPs in response to basal ABA signaling. Further studies are needed to verify this hypothesis.

ZmbZIP75 accumulates predominantly in a phosphorylated state in developing embryo, especially during late kernel development (Figure 3G). ABA significantly enhances ZmbZIP75 phosphorylation, which increases its protein stability and transactivation ability (Figure 8E-8H; Supplemental Figure S19C). ZmSnRK2.10, a member of the subclass III SnRK2 family, phosphorylates ZmbZIP75 at the conserved residue Ser129 (Figure 8D; Supplemental Figure S9A). Mutation of Ser129 to Ala reduces ZmbZIP75 transcriptional activation activity and fails to rescue the ABA-insensitive phenotype in the *abi5* mutant (Supplemental Figure S20), indicating that Ser129 phosphorylation is essential for ABA response. A-group bZIP TFs have been widely reported to mediate ABA signaling and regulate seed maturation, dormancy, germination and stress responses in a phosphorylation-dependent manner (Yu et al. 2015). Specifically, the ABA-activated AtSnRK2.2/2.3/2.6 in Arabidopsis directly phosphorylate ABI5 to control seed development and dormancy (Nakashima et al. 2009). ZmSnRK2.10 is highly homologous to AtSnRK2.6 (Long et al. 2021). These results collectively demonstrate that during the late stages of maize kernel development, high concentrations of zygotic ABA trigger kernel dehydration and maturation via a highly conserved ABA-ZmSnRK2.10-ZmbZIP75 signal transduction pathway (Figure 10M).

### Potential application of ZmbZIP75 for maize breeding

Grain yield and kernel moisture content are currently critical target traits in maize breeding (Xie et al. 2022). However, these two traits have historically exhibited a significant positive correlation (Eta-Ndu and Openshaw 1992; Chai et al. 2017). Early-maturing varieties, characterized by rapid grain filling and kernel dehydration, exhibit lower kernel moisture content at harvest, a desirable trait for mechanical grain harvesting (Li et al. 2021a). Nonetheless, the shorter grain filling duration in early-maturing varieties leads to decreased grain yield (Gasura et al. 2013). To address this trade-off, increasing planting density is a common strategy to enhance yield per unit area in maize cultivation. However, plant density must be maintained within a reasonable range, as excessive density intensifies competition among plants for light, temperature, and water, ultimately hindering maize yield (Van Averbeke and Marais 1992). Additionally, regions with humid climates or insufficient irrigation may not be suitable for high-density planting (Qin et al. 2016; Liu et al. 2017; Luo et al. 2020). Therefore, maximizing yield per plant while minimizing kernel moisture could be an attractive strategy for modern maize breeding. Indeed, a few new varieties with both high yield and low kernel moisture have been recently released (Wang et al. 2019a; Xu et al. 2022). However, the genetic basis for this favorable trait combination remains unknown.

In this study, overexpression of *ZmbZIP75* resulted in significantly increased kernel size and weight, while kernel moisture at harvest was similar to that of the wild-type (Figure 9A, 9B, and 9F). This contrasts previous observations that large-grain varieties exhibited higher kernel moisture content (Barnwal et al. 2012; Li et al. 2021b). Further dynamic analysis revealed that the kernel dehydration rate after PM was significantly higher in the overexpression lines (Figure 9G). Additionally, the grain filling rate in *ZmbZIP75* overexpression lines was significantly higher than that of the wild type (Figure 9E). Unexpectedly, *ZmbZIP75* overexpression lines maintained a similar grain filling duration to the wild-type (Supplemental Figure S22B), which directly contributed to the increase in final yield. Grain filling duration is determined by kernel water relations, and some new varieties exhibit high dehydration rates during grain filling, leading to premature cessation of storage material accumulation and a shorter grain filling duration (Gambín et al. 2007; Wang et al. 2019a). In contrast, the kernel dehydration rate of *ZmbZIP75* overexpression lines during grain filling was comparable to the wild-type (Figure 9G), ensuring normal storage material accumulation, and thereby extending the grain filling duration.

Moreover, F1 hybrids created using a *ZmbZIP75* overexpression line also showed a significant increase in grain yield per plant (Figure 10A-10F). Remarkably, the kernel moisture content was significantly lower than that of wild-type hybrids, suggesting that in the hybrid background, the overexpression of *ZmbZIP75* had a stronger enhancing effect on the kernel dehydration rate (Figure 10I, 10K, and 10L). Importantly, the *ZmbZIP75* overexpression hybrids had no apparent negative effect on other agronomic traits (Supplemental Figure S23). This study collectively demonstrates the significant potential of ZmbZIP75 in breeding high-yield, low-kernel-moisture maize varieties for future maize production.

## Materials and methods

### Plant materials and growth conditions

The maize inbred line B73 was planted in Wen Jiang field in Chengdu, China. We chose plants that grow rapidly at the same rate, were pollinated at the same day and defined as 0 DAP. At 9-DAP, kernels were selected from the middle region of well-pollinated ears, and then incubated in 50 ml of MS medium (3% sucrose) with 0 μM (final concentration), 100 μM or 300 μM of ABA and under 28 °C in the dark for different days. The ABA treatment *in vitro* was performed as described previously (Li et al., 2018b; Seebauer et al., 2018).

The 204 diverse maize inbred lines from tropical and temperate backgrounds are used for association analysis. These lines were kindly provided by Professor Zuxin Zhang of Huazhong Agricultural University, Wuhan, China. More detailed information of these diverse inbred lines was listed in Supplemental Data Set S1. The association population were grown in three different environments: 19CZ (2019 Chongzhou, Sichuan), 19HN (2019 Xinxiang, Henan), and 20CZ (2020 Chongzhou, Sichuan). Well-pollinated ears in the middle of each row were selected to measure kernel agronomic traits, including the 100-kernel weight, kernel length, width and thickness.

The ethyl methanesulfonate (EMS)-mutagenized of *dg1* (EMS4-006287) was obtained from the public EMS mutant library (http://maizeems.qlnu.edu.cn/). To create CRISPR-Cas9 mutants, the 20 bp guide RNA editing sequence targeting the first exon of *ZmbZIP75* was selected and cloned into the CRISPR-Cas9 expression vector (Zhu et al., 2016), the *zmbzip75* CRISPR-Cas9 transgenic lines were generated in the KN5585 background by Wimi Biotechnology (Jiangsu). The obtained knockout lines were backcrossed with KN5585 to generate the BC_1_ seeds. The BC_1_ seeds were planted and screened the heterozygous seedlings without the CRISPR/Cas9 cassette by PCR amplification and sequencing. The obtained plants were grown for self-pollination and further screened the homozygous *zmbzip75* kernels and WT kernels from the same ear through genotyping.

To create overexpression transgenic lines, the *ZmbZIP75* coding sequence was amplified and fusion with 3×Flag sequence, cloned into pCAMBIA1305 vector and driven by the *Ubiquitin* promoter. The *ZmbZIP75* overexpression transgenic lines were generated in the B104 background through *Agrobacterium tumefaciens*-mediated transformation (Raji et al., 2018). The transgenic positive lines were identified by bialaphos (bar) resistance and PCR sequencing, backcrossed thrice to B104 inbred line. The null segregants from the self-pollinated *UbiPro:ZmbZIP75/+* ear were propagated and used as wild-type (WT) when comparing with overexpression lines. All these mutants and overexpression plants were grown in Wen Jiang field in Chengdu, China and the Li Guo field in Ledong, China.

*Nicotiana benthamiana* and *Arabidopsis thaliana* of the Columbia-0 (Col-0) accession were grown in the greenhouse at 23°C and 70% relative humidity with a long-day light cycle (16 h light/8 h dark). *Arabidopsis thaliana* of *abi5* mutant and the Columbia-0 (Col-0) were kindly provided by Dr. Chunzhao Zhao at Zhu lab in Purdue University.

### Genotyping

To identify mutant and transgenic materials, including the maize and *Arabidopsis*, the genomic DNA of leaves were extracted using the cetyl trimethylammonium bromide (CTAB) method (Allen et al., 2006). The partial sequences for Bar and *ZmbZIP75* were amplified from genomic DNA using specific primers to genotype the transgenic materials. Construct-free samples were further analyzed by PCR amplification. The genome-edited sites were genotyped by PCR amplification and sequencing. All primers used in this article were listed in Supplemental Data Set S9.

### Physiological analyses

The starch in maize kernels using I_2_ /KI staining was performed as described previously (Li et al., 2018b). The starch contents were determined using the total starch assay kit (Megazyme, K-TSTA) according to the standard protocol of our laboratory (Hu et al., 2021). At least ten dissected kernels that had been stored at -80 °C were ground together in a chilled mortar and pestle, three individual ears as biological replicates.

The ABA contents were determined by Greensword Creation Technology (Wuhan) using LC-MS/MS system according to a previously described procedure with modification (Fu et al., 2012). Briefly, kernels from three independent ears were collected and ground into fine powder, ABA was extracted from 100 mg of powder using 400 µL 90 % methanol.

### RNA extraction and RT-qPCR

The maize tissues and immature kernels were collected by manual dissection and frozen immediately in liquid nitrogen and stored at -80°C before processing. Total RNA was extracted according to the RNA Extraction Kit (Invitrogen) protocol. A NanoDrop 2000 spectrophotometer was used to detect the RNA quality and concentration. RNA was reverse transcribed using the PrimeScript™ RT Reagent Kit with genomic DNA Eraser (Takara). The quantitative RT-PCRs were performed using the SYBR Green PCR Master Mix Kit (Takara), the 2^−ΔΔCt^ method was employed for relative expression of genes calculation and analysis, maize *β-actin* gene was used as the internal reference. The primers used for qRT-PCR were listed in Supplemental Data Set S9.

### Phylogenetic Analysis

Homologous sequences were identified in the National Center for Biotechnology Information (NCBI) nr (nonredundant protein sequences) database by performing a BLASTp search with ZmbZIP75 protein sequences. Amino acid sequences of the A group bZIP transcription factors in *Zea mays*, *Oryza sativa* and *Arabidopsis thaliana* were downloaded from Gramene protein database (http://www.gramene.org/) with the criteria of identify >50%, score >70. Briefly, amino acid sequences were aligned by ClustalX (v1.83) with default parameters and then constructed a phylogenetic tree using the neighbor joining (NJ) method in MEGA (v5.10). Bootstrapping was performed with 1000 replications for phylogeny testing.

### Polyclonal Antibody Preparation and Protein Gel Blot Analysis

To generate anti-ZmbZIP75 antibody, the full-length *ZmbZIP75* ORF was amplified and cloned into the *Eco*RI/*Sal*I sites of pET-28-SUMO (Invitrogen). The anti-ZmbZIP75 antibodies were produced in rabbits by ABclonal (Wuhan). The protein extraction and immunoblot analysis were performed as described previously (Long et al., 2021). Briefly, the materials were ground into fine powder in liquid nitrogen and resuspended in lysis buffer (100 mM HEPES pH 7.5, 5 mM EDTA pH 8.0, 150 mM NaCl, 1 mM MgCl_2_, 0.1% SDS, 10% Glycerol [v/v], 5 mM DTT, 50 µM PMSF and 1× proteinase inhibitor cocktail) for proteins extraction. Protein concentration was determined using Bio-Rad protein assay and about 15 µg protein fractions were transferred to SDS loading buffer and separated by SDS-PAGE (12.5% acrylamide). Western blotting was performed according to the protocol of our laboratory (Li et al., 2018b). The primary antibodies anti-Flag (ABclonal, AE092), anti-Actin (ABclonal, AC009), anti-Histone H3 (Abmart, T61102), anti-His (Abmart, M20001) and anti-GST (ABclonal, AE001) were purchased. The secondary antibodies to detect the primary antibodies were goat anti-rabbit (ABclonal, AS014) and goat anti-mouse IgG-horseradish peroxidase (HRP) (ABclonal, AS003), respectively.

### Immunofluorescence assays

Immunofluorescence assays was performed as described previously (Yamaji et al., 2017). Briefly, developing B73 kernels at 15-DAP were cut longitudinally with a thickness of about 1 mm, incubated with or without ZmbZIP75 antibody and observed under a confocal laser scanning microscope (LSM880, Carl Zeiss) with an excitation wavelength at 488 nm.

### Subcellular localization

The ORF of ZmbZIP75 without the stop codon was amplified and cloned into the vector pCAMBIA2300. The empty vector pCAMBIA2300 was used as control. These constructs were transformed into onion epidermal cells by particle bombardment. The bombarded onion epidermal cells were photographed after 24 h incubation. The green fluorescent signal was observed by the A1R-si Laser Scanning Confocal Microscope (Nikon, Japan) under blue excitation light at 488 nm. The nucleus was stained by1 µg/mL of 4’,6-diamidino-2-phenylindole (DAPI).

### Transcriptional activation analysis

The ORF and truncated sequences of ZmbZIP75 were amplified and cloned into the vector pGBKT7. The empty vector pGBKT7 and pGBKT7-53 were used as negative control and positive control, respectively. All the constructs were transformed into the yeast strain Y2H by PEG/LiAc method, and the yeast transformants were screened on SD/-Trp-Leu plates and SD/-Trp-Leu-His-Ade plates according to the Yeast Protocols Handbook. In addition, the ORF and truncated sequences of ZmbZIP75 were amplified and cloned into a pBXG1 plasmid with minor modifications. The Luc gene driven by the *35S* mini promoter containing a 5xGal4 binding site was used as a reporter plasmid. The Renilla Luc driven by the *35S* promoter served as the reference vector. The empty vector or fused with VP16 was used as negative control and positive control, respectively. These plasmids were transfected into maize leaves protoplasts and the ratio of Luc/Ren activity were performed as described previously (Hu et al., 2020).

### Phos-Tag assay

To detect the phosphorylation of ZmbZIP75 in kernel, endosperm and embryo at the different developmental stages, total proteins were extracted from kernels and then separated on a 10% (v/w) SDS-PAGE gel with 50 mM of Phos-tag (Wako) and 100 μM MnCl_2_. Immunoblot analysis was performed according to the protocol provided by Wako Chemicals and detected with anti-ZmbZIP75 antibody.

### Phenotypic characterizations of mutant or overexpressed kernels

For the mutants, we screened the homozygous and WT kernels from the self-pollinated heterozygous ears to avoid the influence of potential influence of individual and environmental difference on kernel weight and size. For the overexpression lines, we chose plants that grow rapidly at the same rate, and well-pollinated ears were used for future measurement experiments. Besides, kernels from the middle region of the ears were used for the measurements.

The mature kernels were used to measure kernel length, width, and thickness using a vernier caliper. For the fresh weight and dry weight of kernels at different developmental stages, we collected immature kernels every 3 days starting at 6 DAP and measured the kernel fresh weight (W1) using a precision balance. After the fresh weight was measured, the kernels were dried at 60°C to a constant weight, and the dry weight (W1) was measured. The formulae used for kernel water content (KMC) and kernel dehydration rate (KDR) are as follows:

KMC(%)=[(W1-W2)/W1]×100%; KDR(%)=[(KMC_i_-KMC_i+1_)/(t_i_-t_i+1_)]×100%

W1 is the kernel fresh weight, and W2 is the final weight after drying. *i* is the *ith* measured time, and, *t_i_* is the corresponding day after pollination (9, 12, 15, 18, 21, 24, 27, 30, 33, 36, 39 or 42), KMC_i_ is the corresponding KMC at the corresponding day. The mature kernels were collected for measurement of starch, soluble sugar,

lipids and protein. Total starch contents were measured using the total starch assay kit (Megazyme, K-TSTA), soluble sugar contents were measured using the soluble sugar quantification kit (Comin, KT-1-Y) according to the manufacturer’s instructions, respectively. Lipids, total protein, zein and nonzein protein contents were measured according to a described previously (Yang et al., 2021). In addition, the zein and non-zein protein were loaded on 15% and 12% SDS-PAGE gels and analyzed by Coomassie brilliant blue staining, respectively. The 15-DAP immature kernels were collected for metabolite measurements using the LC-MS/MS system analysis according to a described previously (Yang et al., 2018).

### Histocytochemical Analysis

For light microscopy analysis, the immature kernels at 10,15-DAP and mature kernels were longitudinally cutting and observed via light microscopy. In addition, the immature kernels at 10 and 15-DAP were fixed in FAA buffer for paraffin preparation. The sections were cut by a Leica microtome, stained with hematoxylin eosin and imaged under a bright field by a Leica microscope.

For scanning electron microscopy, the mature kernels were latitudinally cut with a razor blade and photographed with a scanning electron microscopy (Hitachi; S3400N) according to the described previously (Wang et al., 2011).

For transmission electron microscopy, the immature kernels at 15-DAP were sliced and fixed in phosphate buffer (pH 7.2) with 2.5% (w/v) glutaraldehyde. The sections were examined and photographed with a transmission electron microscope (Hitachi; H7600).

### RNA-Seq analysis

For RNA-Seq, total RNA was extracted from 15-DAP kernels of homozygous *zmbzip75* and WT which were harvested from three *zmbzip75/+* self-pollination ears separately. The RNA libraries were constructed and sequenced by APTBIO (Shanghai). After sequencing, raw reads were subjected to quality control by FastQC (v.0.11.9) and mapped to the reference maize genome (B73 RefGen_v4, AGPv4) using Hisat2 (v2.0.5). The gene expression value was estimated by FeatureCounts (v2.0.1) and the differentially expression genes (DEGs) were identified with the thresholds of a *P*-value < 0.01 and a log2FC (fold change) > 1.0. The GO enrichment profile and KEGG enrichment profile of DEGs were performed using the program Phyper (http://www.geneontology.org/) and KEGG Mapper (http://www.genome.jp/kegg/pathway.html), respectively.

### ChIP-Seq and ChIP-qPCR

The experiments were performed as previously described (Li et al., 2015) with minor modifications. Briefly, 15-DAP kernels were dissected and cross-linked in buffer with 1% formaldehyde on ice under vacuum infiltration twice for 15 min. Fixation was stopped by adding 1.25 M glycine, then the fixed sample were washed three times with ddH_2_O and dried with blotting paper. About 3.5g fixed materials were ground into a fine powder used for chromatin isolation, and the isolated chromatin was sonicated to fragments ranging from 200 to 500 bp. The DNA/protein complex was immunoprecipitated by anti-ZmbZIP75 antibody and IgG control, respectively. The precipitated DNA was purified and digested by proteinase K, dissolved in distilled water used for further experiments. For ChIP-Seq, the ChIP-DNA and input DNA libraries were constructed and sequenced by Novogene (Shanghai). For ChIP-qPCR, the immunoprecipitated and input DNA were performed using the SYBR Green PCR Master Mix Kit (Takara) with gene-specific primers. Relative enrichment was represented by input (%). The primers used for ChIP-qPCR were listed in Supplemental Data Set S9.

### Electrophoretic mobility shift assay (EMSA)

Oligonucleotide probes were synthesized and labeled following the protocol of the Pierce Biotin 3′ End DNA Labeling Kit (Thermo scientific). The His-fused ZmbZIP75 was expressed in *Transetta* (DE3) cells and purified with Ni-HA beads (QIAGEN). The recombinant protein was incubated with biotin-labeled probe, competition probe, or mutant probe following the protocol of LightShift Chemiluminescent EMSA kit (Thermo scientific). The following binding signal was visualized using a Chemiluminescent Nucleic Acid Detection Module Kit (Thermo scientific).

### Particle bombardment and transient expression assays

The particle bombardment and transient expression assays were performed according to the protocol of our laboratory (Hu et al., 2012) with minor modifications. Briefly, the maize endosperms at 10-DAP or maize embryos at 18-DAP were detached and cultured in half-strength MS medium for 4h. The *Luc* gene driven by the promoter of target genes was used as reporter vector. ZmbZIP75 or ZmSnRK2.10 driven by the *Ubi* promoter was used as an effector vector. The transient expression vector was modified from the plant expression vector, pBI221. The *Gus* gene driven by *Ubi* promoter or the *RLuc* gene driven by 35s promoter was used as a reference vector. The vectors were co-transformed into the maize endosperms or maize embryos by particle bombardment, then cultured at 28 °C for 24 h in the dark. The ratio of Luc/Gus activity was determined and measured following the dual luciferase reporter assay system (Promega), respectively.

### Protein-protein interaction analysis

For the IP-MS assay, we collected 15-DAP B73 kernels for extracting total protein. The protein G agarose beads (Invitrogen) conjugated with ZmbZIP75 antibody were used for IP. The IP-MS method and LC-MS/MS analysis of immunoprecipitated proteins were performed according to a described previously (Lang et al., 2015). The extracted protein was incubated with anti-ZmbZIP75 antibody and Pierce™ protein A Magnetic Beads (Thermo Scientific; #88802) at 4 °C for 4 h with gentle agitation. IgG was used as the negative control. Next, the beads were washed by the PBS containing 1% Nonidet P-40, finally diluted in Lysis solution (4% SDS; 100mM Tris; 1mM DTT) and boiled for10 min. The samples were analyzed through mass spectrometry detection by Oebiotech (Shanghai).

For the Y2H assay, the ORF of ZmbZIP75 was amplified and cloned into the vector pGADT7, the ORFs of ZmSnRK2s were amplified and cloned into the vector pGBKT7. All the constructs were co-transformed into Y2H gold yeast strain. The Y2H gold yeast cells from each transformation system were screened on SD/-Leu-Trp plates, SD/-Leu-Trp-His, SD/-Leu-Trp-His-Ade and SD/-Ler-Trp-His-Ade +100mM 3AT. The experiment was performed according to the instructions of MacthMaker Gold Y2H system.

For the LCI assay, the ORF of ZmbZIP75 and ZmSnRK2.10 were amplified and fused with cLUC and nLUC, respectively. For the BiFC assay, the ORF of ZmbZIP75 and ZmSnRK2.10 were cloned into pSAT6-C eGFP and pSAT6-N eGFP. Different combinations were transformed into the *Agrobacterium tumefaciens* strain GV3101 and transiently expressed in *N. benthamiana* leaf epidermal cells. The LUC images of *N. benthamiana* leaves were detached after 2 days of cultivation by CCD camera. The green fluorescent signal was observed by the A1R-si Laser Scanning Confocal Microscope (Nikon, Japan) under blue excitation light at 488 nm.

For the Pull-down assay, the recombinant protein GST-ZmSnRK2.10 or GST was purified using Glutathione Sepharose 4B (GE Healthcare) and separately incubated with glutathione beads 4°C for 2h, then equal volumes of His-ZmbZIP75 was added to the reaction and incubated for another 2h. The mixture was washed five times with 1×PBS buffer. The pulled down proteins were transferred to SDS loading buffer and separated by SDS-PAGE (10% acrylamide) and detected by immunoblots using anti-GST antibody and anti-His antibody.

### In vitro Phosphorylation assay

The recombinant protein His-ZmbZIP75, His-ZmbZIP75^48A^ or His-ZmbZIP75^129A^ was incubated with GST-ZmSnRK2.10 in the kinase buffer (50 mM HEPES pH 7.4, 10 mM MgCl2, 1mM DTT, 0.2 mM ATP, and 0.5 uCi of [γ-^32^P]ATP) at 30 °C for 2 h, then the reactions were terminated and separated on a SDS-PAGE (10% acrylamide) gel. The phosphorylated bands were detected by autoradiography (Typhoon 9410 imager).

### Protein stability assay

In vitro cell-free stability assay, total proteins were extracted from the 18-DAP of kernels. Equal amounts of recombinant protein His-ZmbZIP75 and the recombinant protein GST-ZmSnRK2.10 or GST mixed an equal amount of total proteins together, then 1 mM ATP was added and incubated for the indicated time at 30°C, λPPase treatment was applied to dephosphorylate. ZmbZIP75 abundance was determined by immunoblots with anti-His antibody. In vivo cell-free stability assay, the Flag-fused ZmSnRK2.10 was cloned into the vector pBI221. The embryos at 18 DAP were transformed with or without *ZmSnRK2.10-Flag* plasmid DNA by particle bombardment, then cultured in MS medium (with or without ABA) for 24h. Total protein was extracted and ZmbZIP75 abundance was determined by immunoblots with anti-ZmbZIP75 antibody. Anti-actin antibody was used as loading control. The protein intensities were quantified using ImageJ software.

### Statistical analysis

The statistical analysis was performed using a two-sided Student’s *t*-test. The immunoblot results were conducted by tracing out the individual band and quantified using ImageJ software (https://imagej.nih.gov/ij/).

### Accession numbers

Sequence data from this article can be found in the MaizeGDB (https://maizegdb.org) database under the following accession number: *ZmbZIP75* (Zm00001d012296); *Sh1* (Zm00001d045042); *Bt2* (Zm00001d050032); *Sh2* (Zm00001d044129); *Ss1* (Zm00001d045261); *Su2* (Zm00001d037234); *Du1* (Zm00001d000002); *Wx* (Zm00001d045462); *Sbe1* (Zm00001d014844); *Ae* (Zm00001d016684); *ISA1* (Zm00001d049753); *Glb1* (Zm00001d033447); *WRI1* (Zm00001d005016); *VP1* (Zm00001d042396); *NKD2* (Zm00001d026113); *ZmNAC130* (Zm00001d008403); *ZmSnRK2.10* (Zm00001d013736).

## Supplemental Data

**Supplemental Figure S1.** Schematic diagram of maize kernels cultured *in vitro*.

**Supplemental Figure S2.** The EMS mutation of *DG1* in the B73 background.

**Supplemental Figure S3.** Motifs enrichment analysis of the promoters of starch synthesis-related genes which were induced by ABA.

**Supplemental Figure S4.** The expression of *ZmbZIP75* is induced by ABA.

**Supplemental Figure S5.** Expression patterns of Group-A bZIP transcription factors in maize.

**Supplemental Figure S6.** Phylogenetic analysis of ZmbZIP75.

**Supplemental Figure S7.** Immunofluorescence assays of the ZmbZIP75 protein accumulation in 15-DAP kernel.

**Supplemental Figure S8.** Transactivation analysis of ZmbZIP75 in yeast.

**Supplemental Figure S9.** Analysis of phosphorylation residues on ZmbZIP75.

**Supplemental Figure S10.** CRISPR-Cas9-based mutation of Zm*bZIP75* and the phenotypes of *zmbzip75*.

**Supplemental Figure S11.** Phenotypic comparison of WT and *zmbzip75-15#* throughout kernel development.

**Supplemental Figure S12.** Kernel phenotype of the self-pollinated *zmbzip75*-15-/+ developing ears.

**Supplemental Figure S13.** SDS-PAGE analysis of nonzein (left) and zein (right) proteins of wild-type and z*mbzip75* mature kernels.

**Supplemental Figure S14.** Metabolome analysis in wild-type and *zmbzip75-15*.

**Supplemental Figure S15.** qRT-PCR validation of selected genes.

**Supplemental Figure S16.** ZmbZIP75 interacts with VP1 to synergistically enhance the expressions of highly embryo-expressed genes.

**Supplemental Figure S17.** Genome-wide identification of ZmbZIP75 binding sites and target genes by ChIP-Seq.

**Supplemental Figure S18.** ZmbZIP75 directly binds to the promoters of *NKD2* and *NAC130* to activate their transcription.

**Supplemental Figure S19.** ZmSnRK2.10 mediates ABA to enhance the phosphorylation of ZmbZIP75.

**Supplemental Figure S20.** The phosphorylation of ZmbZIP75 at Ser129 is essential for ABA response.

**Supplemental Figure S21.** Generation of *ZmbZIP75*-overexpression lines and phenotype analysis.

**Supplemental Figure S22.** Comparisons of 10-kernel dry weight and kernel moisture content of WT and *ZmbZIP75*-overexpression lines.

**Supplemental Figure S23.** The plant, ear and kernel traits of four hybrids derived from overexpression line OE3 and B104.

**Supplemental Table S1.** The RNA-seq data quality and read mapping summary.

**Supplemental Table S2.** The High-Confidence potential targets of ZmbZIP75.

**Supplemental Data Set S1.** Identification of 16 putative A-group bZIP TFs in the maize reference genome.

**Supplemental Data Set S2.** The information of 204 maize inbred lines.

**Supplemental Data Set S3.** The information of 285 SNPs located in ZmbZIP75.

**Supplemental Data Set S4.** The association between ZmbZIP75 transcription and hundred grain weight (HGW) in 86 maize inbred lines.

**Supplemental Data Set S5.** Statistical analysis of differentially accumulated metabolites in the positive and negative models.

**Supplemental Data Set S6.** The differentially expressed genes (DEGs) identified in zmbzip75 and wild-type kernels at 15 DAP.

**Supplemental Data Set S7.** bZIP75 binding sites identified by chromatin immunoprecipitation sequencing (ChIP-Seq) analysis.

**Supplemental Data Set S8.** The direct target DEGs of bZIP75.

**Supplemental Data Set S9.** Primers used in the study.

## Author Contributions

Conceptualization, Y.L., and Y.H.; Methodology, T.L., Y.W., J.Y., Z.L., C.M., Y.X., and D.Z.; Investigation, Y.L., L.G., H.H, and X.F.; Writing-Original Draft, Y.L., and T.L.; Writing-Review & Editing, Y.L., Y.H., J.Z., and H.L.; Project administration, Y.L., and Y.H.

## Acknowledgments

We thank Dr. Haiyang Wang (South China Agricultural University) and Dr. Jingye Fu (Sichuan Agricultural University) for constructive suggestions and critical revisions of the manuscript. We also thank Dr. Chunyi Zhang from the Institute of Biotechnology of Chinese Academy of Agricultural Sciences and Dr. Xiaoduo Lu from Qilu Normal University for distributing the EMS mutant.

## References

Ali, F., Qanmber, G., Li, F., and Wang, Z. (2022). Updated role of ABA in seed maturation, dormancy, and germination. J. Adv. Res. 35:199–214.

Allen, G.C., Flores-Vergara, M., Krasynanski, S., Kumar, S., and Thompson, W. (2006). A modified protocol for rapid DNA isolation from plant tissues using cetyltrimethylammonium bromide. Nat. Protoc. 1:2320–2325.

Amara, I., Odena, A., Oliveira, E., Moreno, A., Masmoudi, K., Pagès, M., and Goday, A. (2011). Insights into Maize LEA Proteins: From Proteomics to Functional Approaches. Plant Cell Physiol. 53:312–329.

Bailey, T.L., Johnson, J., Grant, C.E., and Noble, W.S. (2015). The MEME suite. Nucleic Acids Res. 43:W39–W49.

Barnwal, P., Kadam, D., and Singh, K. (2012). Influence of moisture content on physical properties of maize. Int. Agrophys. 26.

Bates, P.D., Stymne, S., and Ohlrogge, J. (2013). Biochemical pathways in seed oil synthesis. Curr. Opin. Plant Biol. 16:358–364.

Belanger, F.C., and Kriz, A.L. (1989). Molecular characterization of the major maize embryo globulin encoded by the *Glb1* gene. Plant Physiol. 91:636–643.

Buckner, B., Miguel, P.S., Janick-Buckner, D., and Bennetzen, J.L. (1996). The *yl* gene of maize codes for phytoene synthase. Genetics 143:479–488.

Chai, Z., Wang, K., Guo, Y., Xie, R., Li, L., Ming, B., Hou, P., Liu, C., Chu, Z., and Zhang, W. (2017). Current status of maize mechanical grain harvesting and its relationship with grain moisture content. Scientia Agricultura Sinica 50:2036–2043.

Chen, E., Yu, H., He, J., Peng, D., Zhu, P., Pan, S., Wu, X., Wang, J., Ji, C., and Chao, Z. (2023). The transcription factors ZmNAC128 and ZmNAC130 coordinate with Opaque2 to promote endosperm filling in maize. Plant Cell 35:4066–4090.

Chen, J., Zeng, B., Zhang, M., Xie, S., Wang, G., Hauck, A., and Lai, J. (2014). Dynamic transcriptome landscape of maize embryo and endosperm development. Plant Physiol. 166:252–264.

Cheng, Z.J., Zhao, X.Y., Shao, X.X., Wang, F., Zhou, C., Liu, Y.G., Zhang, Y., and Zhang, X.S. (2014). Abscisic acid regulates early seed development in Arabidopsis by ABI5-mediated transcription of SHORT HYPOCOTYL UNDER BLUE1. Plant Cell 26:1053–1068.

Cutler, S.R., Rodriguez, P.L., Finkelstein, R.R., and Abrams, S.R. (2010). Abscisic acid: emergence of a core signaling network. Annu. Rev. Plant Biol. 61:651–679.

Deng, Y., Wang, J., Zhang, Z., and Wu, Y. (2020). Transactivation of Sus1 and Sus2 by Opaque2 is an essential supplement to sucrose synthase-mediated endosperm filling in maize. Plant Biotechnol. J. 18:1897–1907.

Dröge-Laser, W., Snoek, B.L., Snel, B., and Weiste, C. (2018). The Arabidopsis bZIP transcription factor family—an update. Curr. Opin. Plant Biol. 45:36–49.

Erenstein, O., Jaleta, M., Sonder, K., Mottaleb, K., and Prasanna, B. (2022). Global maize production, consumption and trade: Trends and R&D implications. Food Security 14:1295–1319.

Eta-Ndu, J., and Openshaw, S. (1992). Selection criteria for grain yield and moisture in maize yield trials. Crop Sci. 32:332–335.

Finkelstein, R.R. (1994). Mutations at two new Arabidopsis ABA response loci are similar to the *abi3* mutations. Plant J. 5:765–771.

Finkelstein, R.R., and Lynch, T.J. (2000). The Arabidopsis abscisic acid response gene ABI5 encodes a basic leucine zipper transcription factor. Plant Cell 12:599–609.

Fu, J., Chu, J., Sun, X., Wang, J., and Yan, C. (2012). Simple, Rapid, and Simultaneous Assay of Multiple Carboxyl Containing Phytohormones in Wounded Tomatoes by UPLC-MS/MS Using Single SPE Purification and Isotope Dilution. Anal. Sci. 28:1081–1087.

Furihata, T., Maruyama, K., Fujita, Y., Umezawa, T., Yoshida, R., Shinozaki, K., and Yamaguchi-Shinozaki, K. (2006). Abscisic acid-dependent multisite phosphorylation regulates the activity of a transcription activator AREB1. Proc. Natl. Acad. Sci. 103:1988–1993.

Gambín, B.L., Borrás, L., and Otegui, M.E. (2007). Kernel water relations and duration of grain filling in maize temperate hybrids. Field Crops Res. 101:1–9.

Gao, S., Ming, B., Li, L.-l., Yin, X.-b., Xue, J., Wang, K.-r., Xie, R.-z., and Li, S.-k. (2021). Relationship and distribution of in-field dry-down and equilibrium in maize grain moisture content. Agricultural and Forest Meteorology 304:108409.

Gasura, E., Setimela, P., Edema, R., Gibson, P.T., Okori, P., and Tarekegne, A. (2013). Exploiting grain-filling rate and effective grain-filling duration to improve grain yield of early-maturing maize. Crop Sci. 53:2295–2303.

Gomez, E., Royo, J., Muniz, L.M., Sellam, O., Paul, W., Gerentes, D., Barrero, C., Lopez, M., Perez, P., and Hueros, G. (2009). The maize transcription factor myb-related protein-1 is a key regulator of the differentiation of transfer cells. Plant Cell 21:2022–2035.

Gontarek, B.C., Neelakandan, A.K., Wu, H., and Becraft, P.W. (2016). NKD transcription factors are central regulators of maize endosperm development. Plant Cell 28:2916–2936.

Hannah, L.C., and James, M. (2008). The complexities of starch biosynthesis in cereal endosperms. Curr. Opin. Biotechnol. 19:160–165.

Hattori, T., Totsuka, M., Hobo, T., Kagaya, Y., and Yamamoto-Toyoda, A. (2002). Experimentally determined sequence requirement of ACGT-containing abscisic acid response element. Plant Cell Physiol. 43:136–140.

Hickey, L.T., N. Hafeez, A., Robinson, H., Jackson, S.A., Leal-Bertioli, S.C.M., Tester, M., Gao, C., Godwin, I.D., Hayes, B.J., and Wulff, B.B.H. (2019). Breeding crops to feed 10 billion. Nat. Biotechnol. 37:744–754.

Hu, Y., Li, Y., Zhang, J., Liu, H., Tian, M., and Huang, Y. (2012). Binding of ABI4 to a CACCG motif mediates the ABA-induced expression of the *ZmSSI* gene in maize (*Zea mays* L.) endosperm. J. Exp. Bot. 63:5979–5989.

Hu, Y., Song, D., Gao, L., Ajayo, B.S., Wang, Y., Huang, H., Zhang, J., Liu, H., Liu, Y., and Yu, G. (2020). Optimization of isolation and transfection conditions of maize endosperm protoplasts. Plant Methods 16:1–15.

Hu, Y., Li, Y., Weng, J., Liu, H., Yu, G., Liu, Y., Xiao, Q., Huang, H., Wang, Y., and Wei, B. (2021). Coordinated regulation of starch synthesis in maize endosperm by microRNAs and DNA methylation. Plant J. 105:108–123.

Huang, A. (1996). Oleosins and oil bodies in seeds and other organs. Plant Physiol. 110:1055.

Huang, H., Xie, S., Xiao, Q., Wei, B., Zheng, L., Wang, Y., Cao, Y., Zhang, X., Long, T., and Li, Y. (2016). Sucrose and ABA regulate starch biosynthesis in maize through a novel transcription factor, ZmEREB156. Sci. Rep. 6:27590.

Huang, L., Tan, H., Zhang, C., Li, Q., and Liu, Q. (2021). Starch biosynthesis in cereal endosperms: An updated review over the last decade. Plant Commun. 2.

James, M.G., Denyer, K., and Myers, A.M. (2003). Starch synthesis in the cereal endosperm. Curr. Opin. Plant Biol. 6:215–222.

Johnson, D.R., and Tanner, J. (1972). Calculation of the Rate and Duration of Grain Filling in Corn (*Zea mays* L.) Crop Sci. 12:485–486.

Jones, R.J., and Brenner, M.L. (1987). Distribution of abscisic acid in maize kernel during grain filling. Plant Physiol. 83:905–909.

Kang, B.-H., Xiong, Y., Williams, D.S., Pozueta-Romero, D., and Chourey, P.S. (2009). Miniature1-encoded cell wall invertase is essential for assembly and function of wall-in-growth in the maize endosperm transfer cell. Plant Physiol. 151:1366–1376.

Kang, M., and Zuber, M. (1989). Combining ability for grain moisture, husk moisture, and maturity in maize with yellow and white endosperms. Crop Sci. 29:689–692.

Kato, T., Sakurai, N., and Kuraishi, S. (1993). The changes of endogenous abscisic acid in developing grain of two rice cultivars with different grain size. Jpn. J. Crop Sci. 62:456–461.

Kobayashi, Y., Murata, M., Minami, H., Yamamoto, S., Kagaya, Y., Hobo, T., Yamamoto, A., and Hattori, T. (2005). Abscisic acid-activated SNRK2 protein kinases function in the gene-regulation pathway of ABA signal transduction by phosphorylating ABA response element-binding factors. Plant J. 44:939–949.

Kriz, A.R., Wallace, M.S., and Paiva, R. (1990). Globulin gene expression in embryos of maize viviparous mutants: evidence for regulation of the *Glb1* gene by abscissic acid. Plant Physiol. 92:538–542.

Lang, Z., Lei, M., Wang, X., Tang, K., Miki, D., Zhang, H., Mangrauthia, S.K., Liu, W., Nie, W., and Ma, G. (2015). The methyl-CpG-binding protein MBD7 facilitates active DNA demethylation to limit DNA hyper-methylation and transcriptional gene silencing. Mol. Cell 57:971–983.

Li, C., Yue, Y., Chen, H., Qi, W., and Song, R. (2018a). The ZmbZIP22 transcription factor regulates *27-kD γ-zein* gene transcription during maize endosperm development. Plant Cell 30:2402–2424.

Li, C., Qiao, Z., Qi, W., Wang, Q., Yuan, Y., Yang, X., Tang, Y., Mei, B., Lv, Y., and Zhao, H. (2015). Genome-wide characterization of *cis*-acting DNA targets reveals the transcriptional regulatory framework of *opaque2* in maize. Plant Cell 27:532–545.

Li, L., Ming, B., Gao, S., Wang, K., Hou, P., Jin, X., Chu, Z., Zhang, W., Huang, Z., and Li, H. (2021a). A regional analysis model of maize kernel moisture. Agron. J. 113:1467–1479.

Li, W., Yu, Y., Wang, L., Luo, Y., Peng, Y., Xu, Y., Liu, X., Wu, S., Jian, L., and Xu, J. (2021b). The genetic architecture of the dynamic changes in grain moisture in maize. Plant Biotechnol. J. 19:1195–1205.

Li, Y., Yu, G., Lv, Y., Long, T., Li, P., Hu, Y., Liu, H., Zhang, J., Liu, Y., and Li, W.C.J.P.s.a.i.j.o.e.p.b. (2018b). Combinatorial interaction of two adjacent cis-active promoter regions mediates the synergistic induction of *Bt2* gene by sucrose and ABA in maize endosperm. Plant Sci. 274:332–340.

Lid, S.E., Gruis, D., Jung, R., Lorentzen, J.A., Ananiev, E., Chamberlin, M., Niu, X., Meeley, R., Nichols, S., and Olsen, O.-A. (2002). The *defective kernel 1* (*dek1*) gene required for aleurone cell development in the endosperm of maize grains encodes a membrane protein of the calpain gene superfamily. Proc. Natl. Acad. Sci. 99:5460–5465.

Liu, B., Chen, X., Meng, Q., Yang, H., and van Wart, J. (2017). Estimating maize yield potential and yield gap with agro-climatic zones in China—Distinguish irrigated and rainfed conditions. Agricultural and forest meteorology 239:108–117.

Liu, J., Yu, H., Liu, Y., Deng, S., Liu, Q., Liu, B., and Xu, M. (2020). Genetic dissection of grain water content and dehydration rate related to mechanical harvest in maize. BMC Plant Biol. 20:1–16.

Long, T., Xu, B., Hu, Y., Wang, Y., Mao, C., Wang, Y., Zhang, J., Liu, H., Huang, H., and Liu, Y. (2021). Genome-wide identification of ZmSnRK2 genes and functional analysis of ZmSnRK2. 10 in ABA signaling pathway in maize (*Zea mays* L). BMC Plant Biol. 21:309.

Lu, X., Liu, J., Ren, W., Yang, Q., Chai, Z., Chen, R., Wang, L., Zhao, J., Lang, Z., and Wang, H. (2018). Gene-indexed mutations in maize. Mol. Plant 11:496–504.

Luo, N., Wang, X., Hou, J., Wang, Y., Wang, P., and Meng, Q. (2020). Agronomic optimal plant density for yield improvement in the major maize regions of China. Crop Sci. 60:1580–1590.

Ma, B., Zhang, L., and He, Z. (2023). Understanding the regulation of cereal grain filling: The way forward. Journal of Integrative Plant Biology 65:526–547.

Maiorano, A., Fanchini, D., and Donatelli, M. (2014). MIMYCS. Moisture, a process-based model of moisture content in developing maize kernels. Eur. J. Agron. 59:86–95.

Massigoge, I., Carcedo, A., Lingenfelser, J., Hefley, T., Prasad, P.V., Berning, D., Lira, S., Messina, C.D., Rice, C.W., and Ciampitti, I. (2023). Maize planting date and maturity in the US central Great Plains: Exploring windows for maximizing yields. Eur. J. Agron. 149:126905.

McCarty, D.R., Carson, C.B., Stinard, P.S., and Robertson, D.S. (1989). Molecular Analysis of *viviparous-1*: An Abscisic Acid-Insensitive Mutant of Maize. Plant Cell 1:523–532.

McCarty, D.R., Hattori, T., Carson, C.B., Vasil, V., Lazar, M., and Vasil, I.K. (1991). The Viviparous-1 developmental gene of maize encodes a novel transcriptional activator. Cell 66:895–905.

Miao, C., Xiao, L., Hua, K., Zou, C., Zhao, Y., Bressan, R.A., and Zhu, J.-K. (2018). Mutations in a subfamily of abscisic acid receptor genes promote rice growth and productivity. Proc. Natl. Acad. Sci. 115:6058–6063.

Nakashima, K., Fujita, Y., Kanamori, N., Katagiri, T., Umezawa, T., Kidokoro, S., Maruyama, K., Yoshida, T., Ishiyama, K., and Kobayashi, M. (2009). Three Arabidopsis SnRK2 protein kinases, SRK2D/SnRK2.2, SRK2E/SnRK2.6/OST1 and SRK2I/SnRK2.3, involved in ABA signaling are essential for the control of seed development and dormancy. Plant Cell Physiol. 50:1345–1363.

Neill, S., Horgan, R., and Parry, A. (1986). The carotenoid and abscisic acid content of viviparous kernels and seedlings of *Zea mays* L. Planta 169:87–96.

Niu, L., Du, C., Wang, W., Zhang, M., Wang, W., Liu, H., Zhang, J., and Wu, X. (2022). Transcriptome and co-expression network analyses of key genes and pathways associated with differential abscisic acid accumulation during maize seed maturation. BMC Plant Biol. 22:359.

Ober, E.S., and Setter, T.L. (1992). Water deficit induces abscisic acid accumulation in endosperm of maize viviparous mutants. Plant Physiol. 98:353–356.

Plett, S. (1994). Corn kernel breakage as a function of grain moisture at harvest in a prairie environment. Can. J. Plant Sci. 74:543–544.

Pouvreau, B., Baud, S., Vernoud, V., Morin, V., Py, C., Gendrot, G., Pichon, J. P., Rouster, J., Paul, W., & Rogowsky, P. M. (2011). Duplicate maize Wrinkled1 transcription factors activate target genes involved in seed oil biosynthesis. Plant physiology, 156(2), 674–686.

Prioul, J.L., Méchin, V., Lessard, P., Thévenot, C., Grimmer, M., Chateau-Joubert, S., Coates, S., Hartings, H., Kloiber-Maitz, M., and Murigneux, A. (2008). A joint transcriptomic, proteomic and metabolic analysis of maize endosperm development and starch filling. Plant Biotechnol. J. 6:855–869.

Qin, P., Zhang, G., Hu, B., Wu, J., Chen, W., Ren, Z., Liu, Y., Xie, J., Yuan, H., and Tu, B. (2021). Leaf-derived ABA regulates rice seed development via a transporter-mediated and temperature-sensitive mechanism. Sci. Adv. 7:eabc8873.

Qin, X., Feng, F., Li, Y., Xu, S., Siddique, K.H., and Liao, Y. (2016). Maize yield improvements in China: past trends and future directions. Plant Breeding 135:166–176.

Raji, J.A., Frame, B., Little, D., Santoso, T.J., and Wang, K. (2018). Agrobacterium-and biolistic-mediated transformation of maize B104 inbred. Methods Mol. Biol.:15–40.

Rivin, C.J., and Grudt, T. (1991). Abscisic acid and the developmental regulation of embryo storage proteins in maize. Plant Physiol. 95:358–365.

Seebauer, J.R., Below, F.E.J.M.M., and Protocols (2018). Use of in vitro kernel culture to study maize nitrogen and carbohydrate metabolism. Maize: Methods and Protocols:3–13.

Tao, F., Zhang, S., Zhang, Z., and Rötter, R.P. (2014). Maize growing duration was prolonged across China in the past three decades under the combined effects of temperature, agronomic management, and cultivar shift. Global Change Biol. 20:3686–3699.

Van Averbeke, W., and Marais, J. (1992). Maize response to plant population and soil water supply: I. Yield of grain and total above-ground biomass. South African Journal of Plant and Soil 9:186–192.

Wang, G., Sun, X., Wang, G., Wang, F., Gao, Q., Sun, X., Tang, Y., Chang, C., Lai, J., and Zhu, L. (2011). Opaque7 encodes an acyl-activating enzyme-like protein that affects storage protein synthesis in maize endosperm. Genetics 189:1281.

Wang, X., Wang, X., Xu, C., Tan, W., Wang, P., and Meng, Q. (2019a). Decreased kernel moisture in medium-maturing maize hybrids with high yield for mechanized grain harvest. Crop Sci. 59:2794–2805.

Wang, X., Guo, C., Peng, J., Li, C., Wan, F., Zhang, S., Zhou, Y., Yan, Y., Qi, L., and Sun, K. (2019b). ABRE-BINDING FACTORS play a role in the feedback regulation of ABA signaling by mediating rapid ABA induction of ABA co-receptor genes. New Phytol. 221:341–355.

Wang, Y., Zhang, J., Sun, M., He, C., Yu, K., Zhao, B., Li, R., Li, J., Yang, Z., and Wang, X. (2021). Multi-Omics analyses reveal systemic insights into maize vivipary. Plants 10:2437.

Wilson, G.F., Rhodes, A., and Dickinson, D. (1973). Some Physiological Effects of Viviparous Genes *vp1* and *vp5* on Developing Maize Kernels. Plant Physiol. 52:350–356.

Wu, H., Galli, M., Spears, C. J., Zhan, J., Liu, P., Yadegari, R., Dannenhoffer, J. M., Gallavotti, A., & Becraft, P. W. (2023). NAKED ENDOSPERM1, NAKED ENDOSPERM2, and OPAQUE2 interact to regulate gene networks in maize endosperm development. Plant Cell, 36(1), 19–39.

Wu, X., Gong, F., Yang, L., Hu, X., Tai, F., and Wang, W. (2015). Proteomic analysis reveals differential accumulation of small heat shock proteins and late embryogenesis abundant proteins between ABA-deficient mutant *vp5* seeds and wild-type Vp5 seeds in maize. Front. Plant Sci. 5:801.

Xie, R., Ming, B., Gao, S., Wang, K., Hou, P., and Li, S. (2022). Current state and suggestions for mechanical harvesting of corn in China. J. Integr. Agric 21:892–897.

Xing, G., Li, Y., Yang, M., Li, C., Song, Y., Wang, T., Yu, L., and Shi, Y. (2023). Changes in grain-filling characteristics of single-cross maize hybrids released in China from 1964 to 2014. J. Integr. Agric. 22:691–700.

Xu, C.-C., Zhang, P., Wang, Y.-Y., Ning, L., Tian, B.-J., Liu, X.-W., Pu, W., and Huang, S.-B. (2022). Grain yield and grain moisture associations with leaf, stem and root characteristics in maize. J. Integr. Agric. 21:1941–1951.

Yamaji, N., Takemoto, Y., Miyaji, T., Mitani-Ueno, N., Oshida, K.T.Y., and Ma, J.F. (2017). Reducing phosphorus accumulation in rice grains with an impaired transporter in the node. Nature 541:92.

Yan, H., Pan, X., Jiang, H., and Wu, G. (2009). Comparison of the starch synthesis genes between maize and rice: copies, chromosome location and expression divergence. Theor. Appl. Genet. 119:815–825.

Yang, J., Zhang, J., Wang, Z., Zhu, Q., and Wang, W. (2001). Hormonal changes in the grains of rice subjected to water stress during grain filling. Plant Physiol. 127:315–323.

Yang, J., Zhang, J., Wang, Z., Xu, G., and Zhu, Q. (2004). Activities of key enzymes in sucrose-to-starch conversion in wheat grains subjected to water deficit during grain filling. Plant Physiol. 135:1621–1629.

Yang, J., Fu, M., Ji, C., Huang, Y., and Wu, Y. (2018). Maize oxalyl-CoA decarboxylase1 degrades oxalate and affects the seed metabolome and nutritional quality. Plant Cell 30:2447–2462.

Yang, T., Guo, L., Ji, C., Wang, H., Wang, J., Zheng, X., Xiao, Q., and Wu, Y. (2021). The B3 domain-containing transcription factor ZmABI19 coordinates expression of key factors required for maize seed development and grain filling. Plant Cell 33:104–128.

Yang, T., Wang, H., Guo, L., Wu, X., Xiao, Q., Wang, J., Wang, Q., Ma, G., Wang, W., and Wu, Y. (2022). ABA-induced phosphorylation of basic leucine zipper 29, ABSCISIC ACID INSENSITIVE 19, and Opaque2 by SnRK2. 2 enhances gene transactivation for endosperm filling in maize. Plant Cell 34:1933–1956.

Yoshida, T., Christmann, A., Yamaguchi-Shinozaki, K., Grill, E., and Fernie, A.R. (2019). Revisiting the basal role of ABA–roles outside of stress. Trends Plant Sci. 24:625–635.

Yu, F., Wu, Y., and Xie, Q. (2015). Precise protein post-translational modifications modulate ABI5 activity. Trends Plant Sci. 20:569–575.

Yu, T., Xin, Y., and Liu, P. (2024). Exogenous abscisic acid (ABA) improves the filling process of maize grains at different ear positions by promoting starch accumulation and regulating hormone levels under high planting density. BMC Plant Biol. 24:80.

Yu, Y., Zhang, H., Long, Y., Shu, Y., and Zhai, J. (2022). Plant Public RNA-seq Database: a comprehensive online database for expression analysis of∼ 45 000 plant public RNA-Seq libraries. Plant Biotechnol. J. 20:806.

Yuan, L., Zhou, T., Li, K., Tian, Y., Xu, Y., Zhang, J., and Yang, J. (2023). Moderate soil drying improves physiological performances and kernel yield of maize. Food and Energy Security 12:e444.

Yuan, Y., Huo, Q., Zhang, Z., Wang, Q., Wang, J., Chang, S., Cai, P., Song, K.M., Galbraith, D.W., and Zhang, W. (2024). Decoding the gene regulatory network of endosperm differentiation in maize. Nat. Commun. 15:34.

Zhang, H., Li, H., Yuan, L., Wang, Z., Yang, J., and Zhang, J. (2012). Post-anthesis alternate wetting and moderate soil drying enhances activities of key enzymes in sucrose-to-starch conversion in inferior spikelets of rice. J. Exp. Bot. 63:215–227.

Zhang, H., Gou, X., Ma, L., Zhang, X., Qu, J., Wang, X., Huang, W., Yan, S., Zhang, X., and Xue, J. (2024). Reveal the kernel dehydration mechanisms in maize based on proteomic and metabolomic analysis. BMC Plant Biol. 24:15.

Zhang, L., Li, X., Gao, Z., Shen, S., Liang, X., Zhao, X., Lin, S., and Zhou, S. (2017a). Regulation of maize kernel weight and carbohydrate metabolism by abscisic acid applied at the early and middle post-pollination stages in vitro. J. Plant Physiol. 216:1–10.

Zhang, L., Liang, X., Shen, S., Yin, H., Zhou, L., Gao, Z., Lv, X., and Zhou, S. (2018). Increasing the abscisic acid level in maize grains induces precocious maturation by accelerating grain filling and dehydration. Plant Growth Regul. 86:65–79.

Zhang, X., and Kaeppler, S. (2017b). Natural variations in maize kernel size: A resource for discovering biological mechanisms. Maize kernel development:204–216.

Zhang, Y., Sun, Q., Zhang, C., Hao, G., Wang, C., Dirk, L.M., Downie, A.B., and Zhao, T. (2019a). Maize VIVIPAROUS1 interacts with ABA INSENSITIVE5 to regulate GALACTINOL SYNTHASE2 expression controlling seed raffinose accumulation. J. Agric. Food Chem. 67:4214–4223.

Zhang, Z., Yang, J., and Wu, Y. (2015). Transcriptional regulation of zein gene expression in maize through the additive and synergistic action of opaque2, prolamine-box binding factor, and O2 heterodimerizing proteins. Plant Cell 27:1162–1172.

Zhang, Z., Zheng, X., Yang, J., Messing, J., and Wu, Y. (2016). Maize endosperm-specific transcription factors O2 and PBF network the regulation of protein and starch synthesis. Proc. Natl. Acad. Sci. 113:10842–10847.

Zhang, Z., Dong, J., Ji, C., Wu, Y., and Messing, J. (2019b). NAC-type transcription factors regulate accumulation of starch and protein in maize seeds. Proc. Natl. Acad. Sci. 116:11223–11228.

Zheng, X., Li, Q., Li, C., An, D., Xiao, Q., Wang, W., and Wu, Y. (2019). Intra-kernel reallocation of proteins in maize depends on VP1-mediated scutellum development and nutrient assimilation. Plant Cell 31:2613–2635.

Zhu, G., Ye, N., Yang, J., Peng, X., and Zhang, J. (2011). Regulation of expression of starch synthesis genes by ethylene and ABA in relation to the development of rice inferior and superior spikelets. J. Exp. Bot. 62:3907–3916.

Zhu, J., Song, N., Sun, S., Yang, W., Zhao, H., Song, W., and Lai, J. (2016). Efficiency and Inheritance of Targeted Mutagenesis in Maize Using CRISPR-Cas9. J. Genet. Genomics 43:25–36.

Zinsmeister, J., Lalanne, D., Terrasson, E., Chatelain, E., Vandecasteele, C., Vu, B.L., Dubois-Laurent, C., Geoffriau, E., Signor, C.L., and Dalmais, M. (2016). ABI5 is a regulator of seed maturation and longevity in legumes. Plant Cell 28:2735–2754.

